# No need for fat: Nutritional preferences affect pollen foraging networks in alpine pollinator communities

**DOI:** 10.64898/2026.05.22.727106

**Authors:** Marielle C. Schleifer, Johann Neumayer, Fabian A. Ruedenauer, Laura Castiglioni, Alexander Keller, Johannes Spaethe, Sara D. Leonhardt

## Abstract

Most flowering plant species rely on insect pollinators for successful cross-pollination. While floral color, scent, and morphology help attract visitors, the primary motivation for visiting a flower is the nutritional reward – nectar and, especially, pollen. Pollen provides a complete set of macro- and micronutrients essential for adult and larval provisioning in many insect flower visitors. However, pollen nutrient composition (i.e., quality) varies widely among plant species, and different visitors may have distinct nutritional requirements. Whether these differences are linked to the selection of plants visited remains poorly understood.

In this study, we investigated whether the nutrient composition of pollen is linked to the visitation patterns of two main alpine pollinator groups: bumblebees and hoverflies. We observed flower visits in the field and identified the origin of pollen collected by bumblebees and hoverflies via metabarcoding. In addition, we analyzed the nutrient profiles of alpine flowering plants from the same habitat, including amino acids, fatty acids, and sterols. We hypothesized that both bumblebees and hoverflies (i) differ in the plant spectrum they were observed on/collected from and visited for pollen collection, (ii) and that preferred nutrient profiles differ between the two pollinator groups. We additionally expected (iii) that, in the studied alpine plant communities, the nutrient composition of pollen of plant species collected by flower visitors is more strongly associated with pollinator identity than with plant phylogeny due to competition for pollinators.

Our results revealed that alpine bumblebees and hoverflies were more similar in their pollen hosts and nutritional preferences than expected, but, for both groups, the network based on observed flower visits differed significantly from that based on pollen collection. Pollen fatty acids and amino acids content was positively correlated, and both bumblebees and hoverflies preferred pollen with low fatty and amino acid content. Pollen sterols did not differ between plants collected by pollinators and non-collected plants. Neither fatty acid, amino acid, nor sterol composition was linked to plant phylogeny. Our findings suggest that not only amino acid content, but also fatty acid content, plays a key role in shaping pollen-collection patterns of flower visitors in alpine ecosystems.

## INTRODUCTION

Animal pollinators are responsible for the successful cross-pollination of more than 80% of wild flowering plant species (Ollerton et al., 2011). The vast majority of these pollinators are insects (Aslan et al., 2013), including Hymenoptera, Lepidoptera, Diptera, and Coleoptera. This mutualistic relationship, in which plants provide resources such as food, shelter or mating sites (Armbruster, 2011) in exchange for pollen transfer, is among the most ecologically significant relationships in nature (Patiny, 2011; van der Kooi et al., 2021). To locate and evaluate flowers, insect pollinators use a set of different floral cues or signals, with color, shape, and scent considered most important (Erickson et al., 2022; Russell et al., 2018; van der Kooi et al., 2023; Whitney & Glover, 2007).

Among the floral rewards, nectar is the primary energy source for most flower visitors (Haslett, 1989; Woodcock et al., 2014). However, nectar contains only low amounts of additional nutrients, such as amino acids (AA), fatty acids (FA), or micronutrients (Leonhardt et al., 2024; Nicolson, 2011). To obtain these additional and essential nutrients, many flower visitors that are restricted to floral resources as source of nutrients feed on pollen to ensure an adequate supply of AAs, FAs, sterols, vitamins, and minerals, which are crucial for rearing brood (e.g., bees: (Vaudo et al., 2015)) and for the development of reproductive organs (e.g., female hoverflies: (Schneider, 1948)). However, the nutrient composition of pollen (hereinafter referred to as pollen quality) strongly varies between different plant species (Crone & Grozinger, 2021; Leonhardt et al., 2024; Roulston et al., 2000; Ruedenauer et al., 2019a; Vaudo et al., 2015). In fact, the quality and diversity of pollen strongly affect bee physiology (Di Pasquale et al., 2013) and can determine pollinator health and fitness (Leonhardt et al., 2024; Parreno et al., 2022). For example, laboratory experiments have demonstrated that bees (i.e., bumblebees and honeybees) are sensitive to varying nutritional compositions and exhibit nutrient-specific perception and regulation (Nebauer et al., 2023; Ruedenauer et al., 2021; Ruedenauer et al., 2019a; Ruedenauer et al., 2020). While elevated amino acid or sterol concentrations in the food do not appear to affect intake or survival rates, even modest increases in FA concentrations can result in strong intake regulation and high mortality (Nebauer et al., 2023; Ruedenauer et al., 2020; Schleifer et al., 2024). Moreover, different pollinators or flower visitors may differ in their nutritional requirements, as shown for different bee species (Barraud et al., 2022), but the precise nutritional requirements, particularly for non-bee pollinators such as hoverflies, still remain largely unknown (Jones & Rader, 2022).

The species-rich alpine plant communities harbor a large spectrum of flowering herbaceous species, most of which depend on insect pollinators for cross-pollination (Körner, 1999). Alpine ecosystems are characterized by extreme climatic conditions, including low temperatures, high winds, and significant exposure to ultraviolet radiation (Inouye, 2020). Moreover, alpine ecosystems experience short growing seasons (Billings, 1973), which may result in competition for pollinators among plant species. This might have led to the adaptation of the plants’ nutritional quality by adjusting the nutritional composition of rewards to the nutritional requirements of their main pollinators in order to increase their attractiveness and thus competitiveness (Ruedenauer et al., 2019b). In these habitats, bumblebees and hoverflies are among the most important pollinators (Inouye, 2020; Warren et al., 1988). Bumblebees and hoverflies exhibit contrasting life-history traits that directly affect resource use and interaction patterns. Bumblebees are eusocial insects with annual colonies, in which workers repeatedly forage and actively transport pollen to provision larvae, whereas solitary hoverflies consume pollen exclusively for their own nutritional needs, particularly for ovarian development (Schneider, 1948; Westrich, 2018). Because larval provisioning is decoupled from adult foraging in hoverflies, their flower visitation is not constrained by a central nesting site, in contrast to bumblebees, which are central-place foragers (Doyle et al., 2020). Alpine communities consequently represent a valuable system to investigate the role of reward nutrition in structuring interaction networks between plants and the two most important flower visitors.

Within this system, our study focuses on three main questions. First, we asked whether alpine bumblebees and hoverflies collect pollen from the same plant species. Second, we examined whether the plant species visited by both pollinator groups differ in their pollen quality as compared to pollen of plants not visited by any of the two pollinator groups. Third, we investigated whether variation in pollen nutrient composition among plant species is shaped by pollen-collection patterns or by plant phylogeny. Based on differences in their life-history traits, we hypothesized that bumblebees and hoverflies would forage on different sets of plant species. As different flower visitors prefer different nutrients (Vaudo et al., 2015), we further expected that nutrient composition of pollen differs between plants visited by bumblebees and those visited by hoverflies. Finally, since FAs were strongly regulated in previously conducted feeding experiments with bees, we hypothesized that FA composition would be more strongly associated with pollen collection patterns than with phylogeny, whereas sterol and AA profiles would show a stronger phylogenetic signal (due to relaxed selection by pollinators) (Mondal & Mandal, 2009; Zu et al., 2021).

## MATERIAL AND METHODS

### Study region

All data were collected in the Hohe Tauern National Park, Austria, which is the country’s largest national park, renowned for its stunning alpine landscapes and rich biodiversity (Bauch & Lainer, 2014). Spanning over 1,800 square kilometers, the park provides a sanctuary for flora and fauna, including rare and endangered plant and animal species, making it a vital area for conservation. Along the Großglockner Hochalpenstraße (a famous mountain road, connecting Bruck in the Salzburg region with Heiligenblut in Carinthia), we observed plant-pollinator interactions and collected flowering plants for pollen nutrient analysis and bumblebees and hoverflies for pollen loads between 1400 and 2500 m a.s.l.

### Observations on flowers and collection of pollen loads

During the summer season of 2023 (June-September), alpine bumblebees and hoverflies were observed visiting flowers and collected in four sampling rounds to analyze their pollen loads. Sampling took place at 20 locations along the Großglockner Hochalpenstraße, where bumblebees and hoverflies were captured using insect nets while visiting flowers. For each flower visitor collected the corresponding plant species was noted. Bumblebees were identified in the field using an identification key (Gokcezade et al., 2010). To facilitate pollen removal from the corbiculae, bees were anesthetized using CO₂ applied via a reusable whipped cream dispenser (Hendi Food Service Equipment, De Klomp, Netherlands). Pollen loads were collected using pointed forceps, which were cleaned with 10% bleach after each use. After pollen removal, the bumblebees regained consciousness after a few minutes and were released. The collected pollen loads were pooled for each sampling round and bumblebee species and stored at -20°C for further analysis. Hoverflies were also observed and collected at the same sites. Since hoverflies do not carry pollen externally but ingest it, they were caught, stored in small vials, and killed in a freezer as soon as possible. After identification in the lab using an identification key (Bothe, 1989), the gastrointestinal system was removed from each hoverfly and pooled for each species and sampling round. We included only those pooled pollen loads of bumblebees and hoverflies in the analysis that comprised at least three individuals per species and sample period. All samples were subsequently stored at -20°C until sent for metabarcoding analysis.

### Collection of flowers and pollen extraction

From 2021 to 2023, between 50 and 500 flower buds were collected from 85 flowering plant species during the summer season for any flowering species found in the study area. After thorough drying, the pollen was extracted using analytical sieves (150µm and 180µm mesh size, Retsch, Haan, Germany) attached to a vortex mixer. This method effectively removed the pollen from the flower buds, while particles larger than 150µm were retained in the sieve. To further purify the pollen samples and remove minor contaminations, a plastic card was rubbed against a paper towel to generate a static charge. The charged card was held above the sample, attracting and removing lightweight contaminants without affecting the pollen grains. The cleaned pollen samples were stored at -20°C until further analysis.

### Pollen metabarcoding

DNA was then extracted from these samples using the Macherey Nagel Food kit (Keller et al., 2015). DNA amounts were normalized between samples, quality controlled and plant taxa identified using Illumina-based ITS2 pollen DNA metabarcoding according to the established laboratory protocol of Sickel et al. (2015) and Campos et al. (2021). Data were processed using the pipeline available at https://github.com/chiras/metabarcoding_pipeline (Leonhardt et al., 2022). Raw reads were merged, quality filtered (200 to 500 bp, no ambiguous bases, ee < 1.0), dereplicated, denoised, and chimaeras removed to generate amplicon sequence variants (ASVs) using VSEARCH v2.15.2 (Rognes et al., 2016). Anomalous ASVs were identified with MetAnoDe v8b35228 (Keller, 2025) and removed. Remaining ASVs were classified at species level using VSEARCH with a ≥97% similarity threshold (top hit) against a localized reference database built with BCdatabaser (Keller et al., 2020). Unclassified reads were subsequently queried against a curated global database (Quaresma et al., 2024), and remaining sequences were assigned using SINTAX (Edgar, 2016) with a confidence threshold of 0.8 against the same database. ASVs not classified at the species level were post-clustered at 98% sequence identity to enable inclusion of unresolved taxa in diversity estimates. Each DNA-based pollen identification was cross-validated using up to two independent databases (NCBI and GBIF). Additionally, all identified plant species were assessed for temporal and spatial plausibility based on their known distributions and flowering periods (Fischer et al., 2008), as well as personal field observations.

### Pollen nutrient analysis

Chemical analyses of pollen nutrients were conducted during winter 2023/2024, focusing on amino acids (AA), fatty acids (FA), and sterols. Free and protein-bound AA were analyzed by ion-exchange chromatography (IEC) following the method of Leonhardt and Blüthgen (2011). Pollen FA content and composition were determined via gas chromatography-mass spectrometry (GC-MS) based on Villagómez et al. (2023). The sterol content and composition were analyzed using GC-MS according to the method adapted from Vanderplanck et al. (2011). For more details, see supplementary materials (S1, S2, S3). The sum of all AAs, FAs, and sterols was used to analyze the total content of each of the nutrients.

### Statistical analysis

All statistical analyses were performed using R (v. 4.4.2.) (R Core Team, 2024). Prior to analysis, data were tested for normality and homogeneity of variances using the Shapiro-Wilk test and Levene’s test, respectively. None of the variables met the assumption of normality, and neither log nor square-root transformation improved the distributional properties. We therefore applied non-parametric tests. Before the statistical analyses are outlined, the replication statement is provided in Table 1.

**Table 1:**
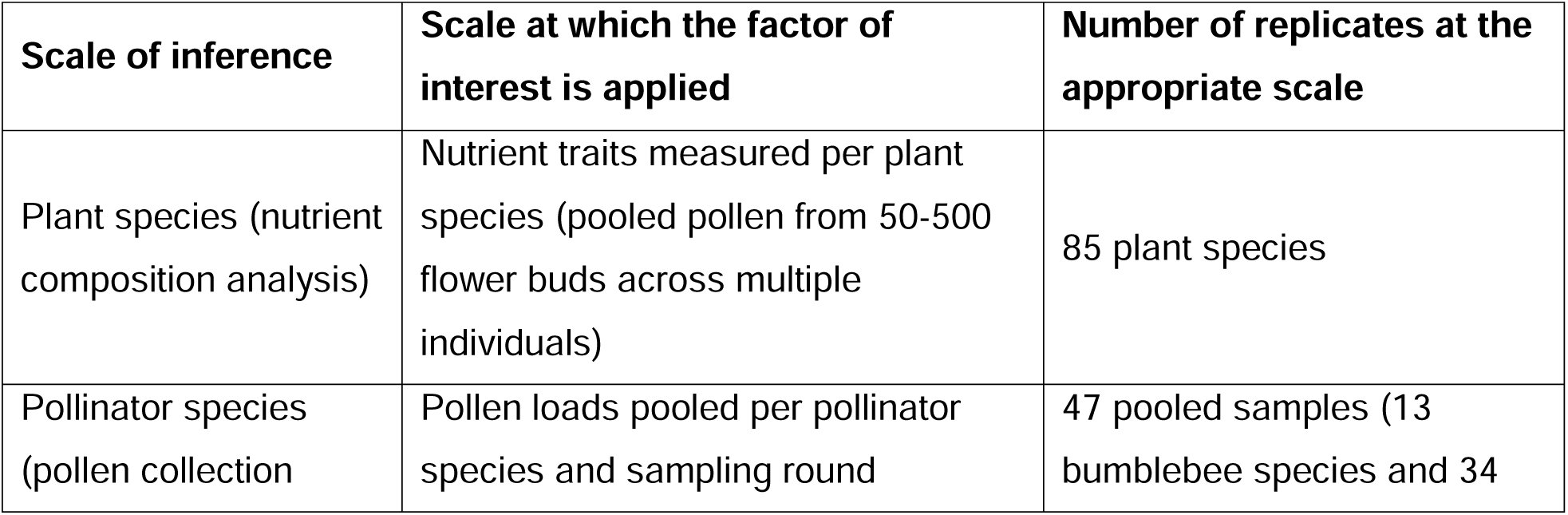
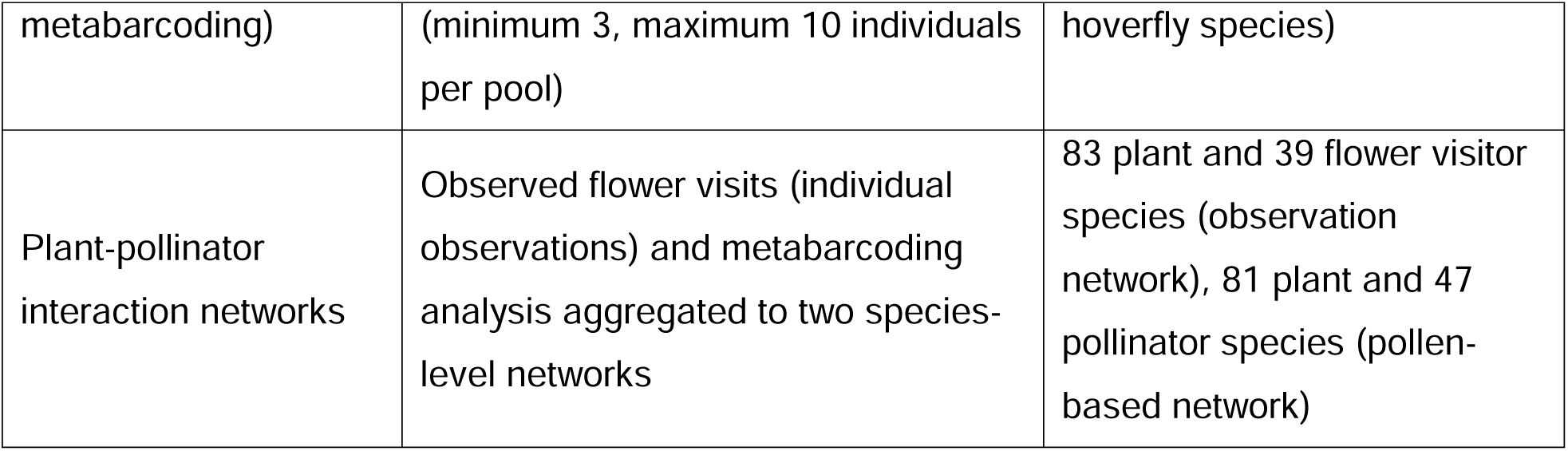
Replication statement.

| Scale of inference | Scale at which the factor of interest is applied | Number of replicates at the appropriate scale |
| --- | --- | --- |
| Plant species (nutrient composition analysis) | Nutrient traits measured per plant species (pooled pollen from 50-500 flower buds across multiple individuals) | 85 plant species |
| Pollinator species (pollen collection) | Pollen loads pooled per pollinator species and sampling round | 47 pooled samples (13 bumblebee species and 34 |
| metabarcoding) | (minimum 3, maximum 10 individuals per pool) | hoverfly species) |
| Plant-pollinator interaction networks | Observed flower visits (individual observations) and metabarcoding analysis aggregated to two species-level networks | 83 plant and 39 flower visitor species (observation network), 81 plant and 47 pollinator species (pollen-based network) |

To visualize the interactions between flower visitors and visited plant species, we constructed quantitative bipartite networks using the *bipartite* package (Dormann et al., 2009). Two separate networks were generated: one based on observed flower visits and a second on metabarcoding data. For each dataset, interaction data were aggregated into a species-by-species matrix, with plant species as lower-level nodes and flower visitor species as upper-level nodes. Network metrics such as connectance, H2’, and d’ were calculated to characterize and compare the overall structure and degree of specialization for both networks. To quantify species-level specialization, d’ was used only for the upper level (flower visitors) in each network. To compare overall specialization across networks, a Mann-Whitney U test was used for all upper-level species, and the effect size was quantified using Cliff’s delta.

Because flowers visited for pollen collection by hoverflies almost completely overlapped with those visited by bumblebees, we did not test for differences between the two pollinator guilds but only tested whether the nutrient composition and content of pollen collected by hoverflies and bumblebees differed from pollen of plants that were not collected by any of the two pollinator groups but flowered at the same time and location. Differences in the nutritional composition between the two plant groups were assessed using non-metric multidimensional scaling (NMDS), followed by PERMANOVA using the *vegan* package (Oksanen et al., 2013). Homogeneity of multivariate dispersions was tested using PERMDISP (also in *vegan*) prior to running PERMANOVA. The different nutrient groups – AAs, FAs, and sterols- were analyzed separately. For FAs, significant differences were found between the groups (collected and non-collected plants), so we also conducted environmental fitting using envfit (*vegan* package). We only show those FAs that were highly significant in the results. Because AA showed a strong horseshoe pattern, the data were arcsine-square-root transformed, which yielded a better result. To check if pollen FAs and AAs concentrations were correlated across plant species, a Spearman correlation was performed. To test for differences in the total concentration of FAs, AAs and the protein:lipid (P:L) ratio between the two plant groups (pollen collected and non-collected), we used Wilcoxon Rank Sum tests.

To finally assess whether pollen nutrient composition was linked to plant phylogeny, we calculated Blomberg’s *K* for each individual nutritional group (FAs, AAs, and sterols) using the package *phytools* (Revell, 2024). The hierarchical taxonomic trees were obtained from phyloT, which constructs tree topologies based on the NCBI taxonomy or the Genome Taxonomy Database (GTDb) (https://phylot.biobyte.de/). Because the tree lacked branch lengths, they were calculated using the Grafen method (Grafen, 1989) implemented in the *ape* package (Paradis et al., 2019).

To account for multiple testing across all nutrient variables, *p*-values were adjusted using the Bonferroni correction.

## RESULTS

### Interaction networks

The two networks, i.e., the one based on observed flower visits (hereafter referred to as the observation network) and the one based on pollen metabarcoding (hereafter referred to as the pollen-based network), revealed distinct interaction patterns and plant compositions (Figure 1 and Figure 2). The observation network included 83 plant and 39 pollinator species (12 bumblebee and 27 hoverfly species) (Figure 1). The pollen-based network contained 81 plant species and 47 flower visitors (13 bumblebee and 34 hoverfly species) (Figure 2). The number of species present in both networks was 32 and 28 for flower visitors and plant species, respectively. The observation network showed low connectance (0.081), with plants visited by an average of three species, and pollinators visiting an average of seven plant species. The pollen-based network was comparatively more connected (0.149). On average, plants interacted with seven insect species and flower visitors with twelve plant species. Network specialization was moderate for the observation network (H2’ = 0.473) and higher for the pollen-based network (H2’ = 0.634). When comparing all upper-level species (i.e., d’ of flower visitors) across networks, specialization was higher in the observation network than in the pollen-based network (U = 90, p < 0.001, effects δ = -0.90).

**Figure 1:**
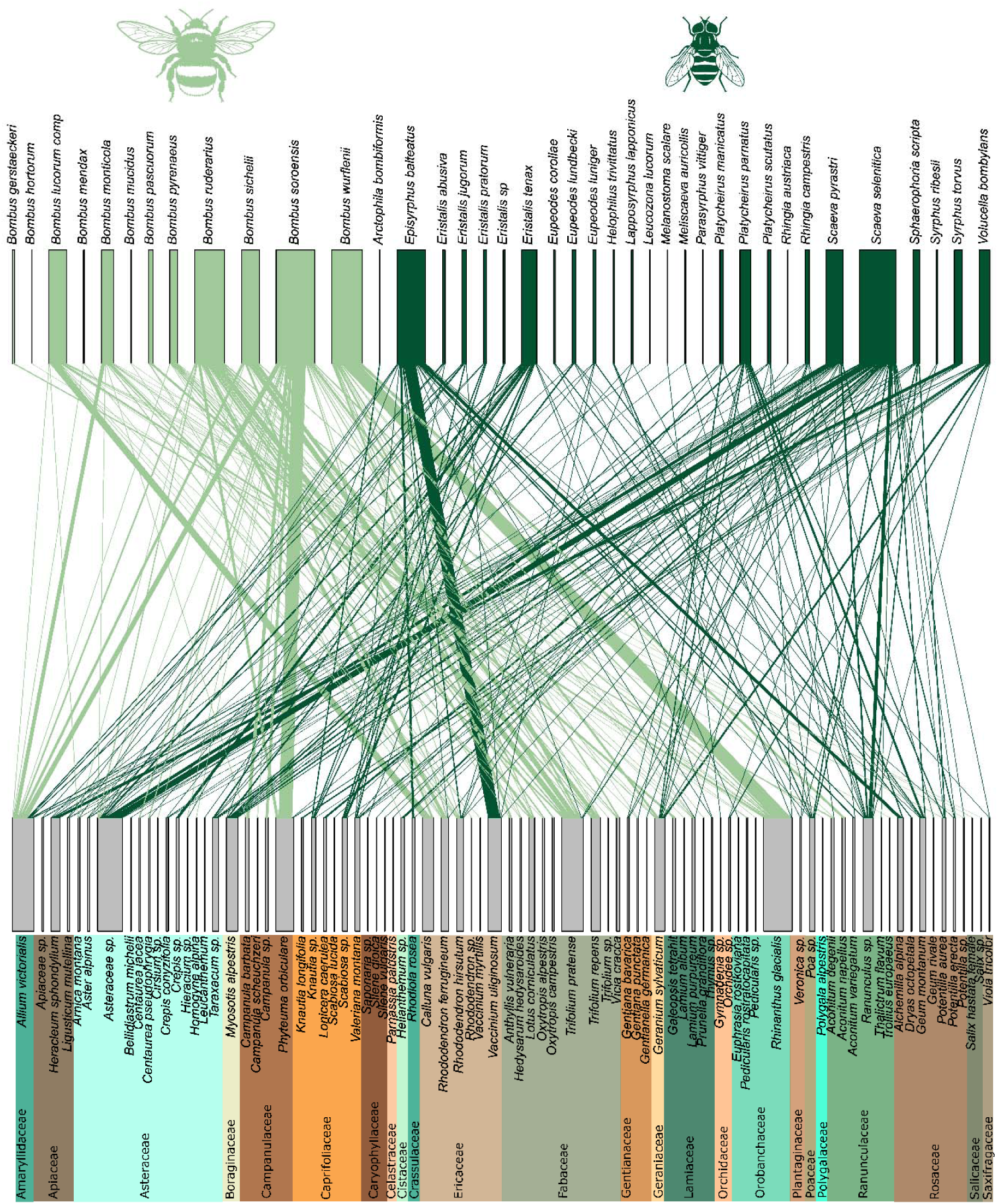
Network based on observation data. Bumblebee species (N = 12) are shown in light green, and hoverfly species (N = 27) in dark green. Plant species (N = 83) are shown in the lower level of the network. Below the plant species, plant families are color-coded and listed underneath. The thickness of the connection lines is proportional to the interaction strength.

**Figure 2:**
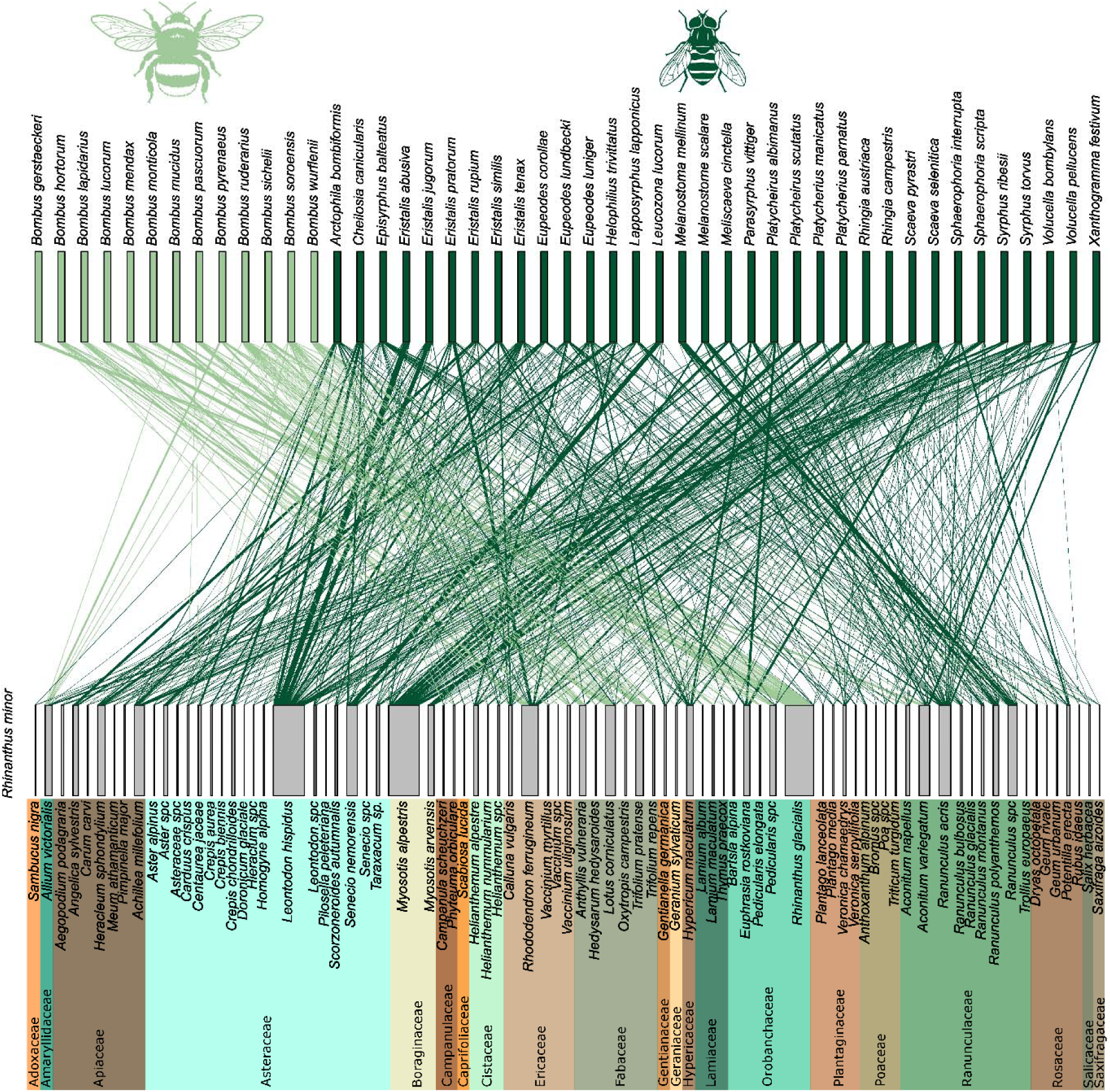
Network based on DNA metabarcoding data. Bumblebee species (N = 13) are shown in light green, and hoverfly species (N = 34) in dark green. Plant species (N = 81) are shown in the lower level of the network. Below the plant species, plant families are color-coded and listed underneath. The thickness of the connection lines is proportional to the interaction strength.

### Pollen nutrient composition

Nutrient profiles of 83 flowering plant species were analyzed (Table S1). For some plant species, all three nutrient groups—AAs, FAs, and sterols—were quantified, while for others, only one or two groups could be quantified due to limited amounts of pollen. All nutrient groups exhibit strong interspecific variation in both total nutrient content and the composition of individual compounds. Total AA concentration varied from 7.55 mg/g pollen (*Myosotis arvensis*) to 475 mg/g pollen (*Ranunculus acris,* Figure 3). Overall, 13 AAs (i.e., aspartic acid, glutamic acid, leucine, lysine, glycine, proline, arginine, phenylalanine, threonine, serine, tyrosine, isoleucine, and histidine) were found in all samples (N = 75). The most abundant AAs were glycine (59.4 mg/g), aspartic acid (53.1 mg/g), and alanine (53.1 mg/g), all found in *Ranunculus acris*. Concentrations of the same AA varied substantially between plant species, i.e., roughly 100-fold (glycine and alanine) to 1000-fold (aspartic acid) between the lowest and highest measured concentrations.

**Figure 3:**
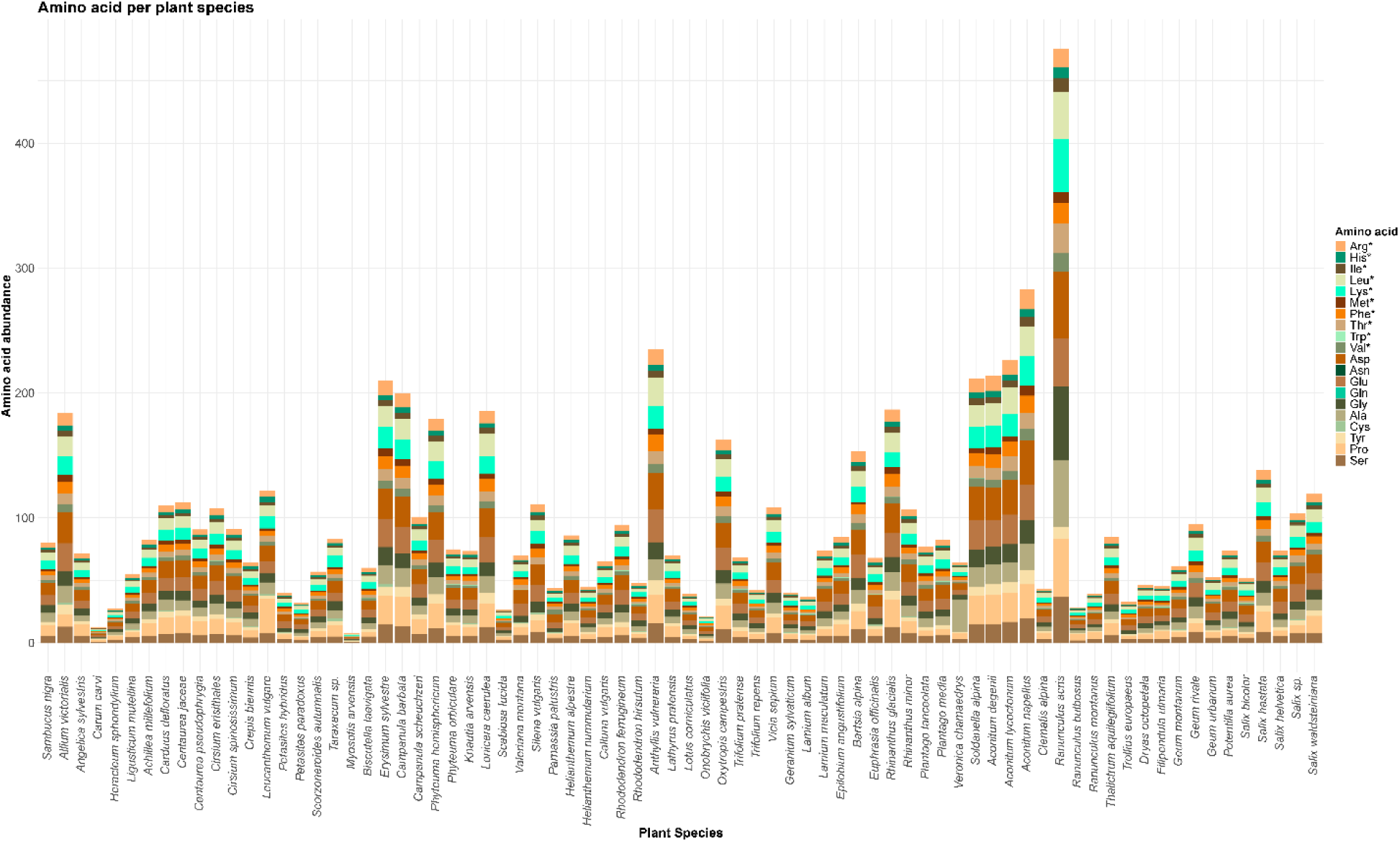
Composition of amino acids (AAs)s in mg/g pollen of 75 plant species. AAs are color-coded. An asterisk after the name indicates essential AAs (EAAs).

Total FA concentration differed from 0.47 mg/g pollen (*Bartsia alpina*) to 412.07 mg/g pollen (*Centaurea pseudophrygia*, Figure 4). Only one FA (octadecadienoic/octadecatrienoic acid) was found in all samples (N = 77); however, linolenic/oleic acid was found in 76 samples, and palmitic acid in 75. Linolenic/oleic acid (127.6 mg/g in *Campanula barbata*), linoleic acid (121.5 mg/g in *Centaurea pseudophrygia*), and palmitic acid (79.3 mg/g in *Centaurea pseudophrygia*) were most abundant. Like AAs, their concentrations also showed pronounced variation across plant species, with the same FA differing approximately 80 (palmitic acid) to 1000 (linoleic acid) times between the lowest and highest concentrations measured.

**Figure 4:**
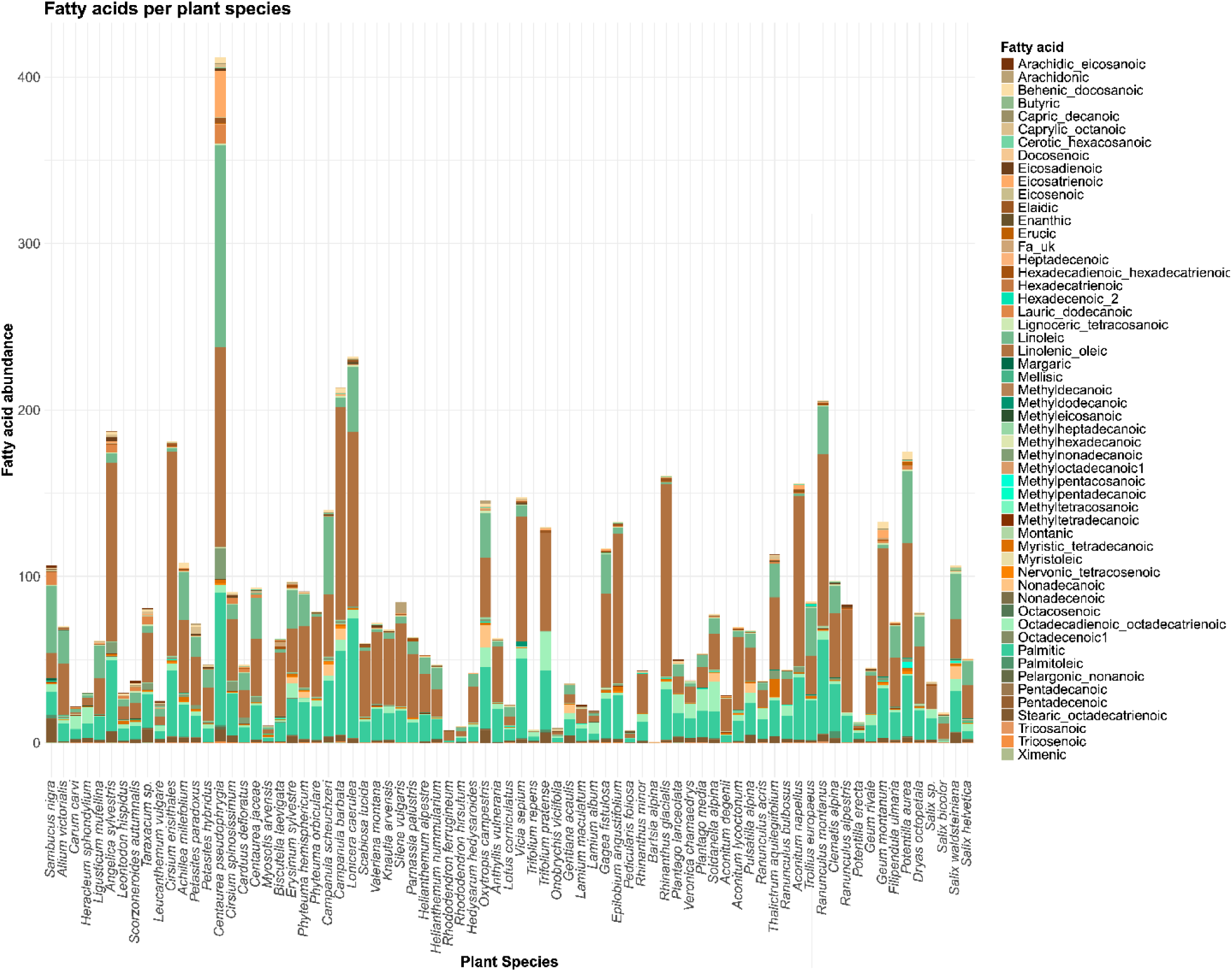
Composition of FAs in mg/g pollen of 77 plant species. FAs are color-coded. Linoleic and linolenic acids are considered essential FAs (EFAs).

Total sterol content varied between 1.07 µg/g pollen (*Myosotis arvensis*) and 40.7 µg/g pollen (*Salix* sp., Figure 5). No sterol was found in all samples (N = 54). The three most common sterols were taraxasterol/sitosterol (found in 51 samples) and cycloartenol/stigmastenol/lanasterol, and delta-5-avenasterol/stigmasterol (found in 50 samples). Cholesterol was found in 48 samples. The most abundant sterols were methylene-cholesterol (18.65 µg/g in *Salix* sp.), delta-5-avenasterol/stigmastanol (16.22 µg/g in *Cirsium eristhales*) and desmosterol (11.88 µg/g in *Geum rivale*). For each of these compounds, concentrations varied widely between plant species, with differences of 100-10,000-fold (methylene-cholesterol) between minimum and maximum concentrations. More information about relative nutrient compositions is provided in the Supplementary Material (Figure S1,Figure S2,Figure S3).

**Figure 5:**
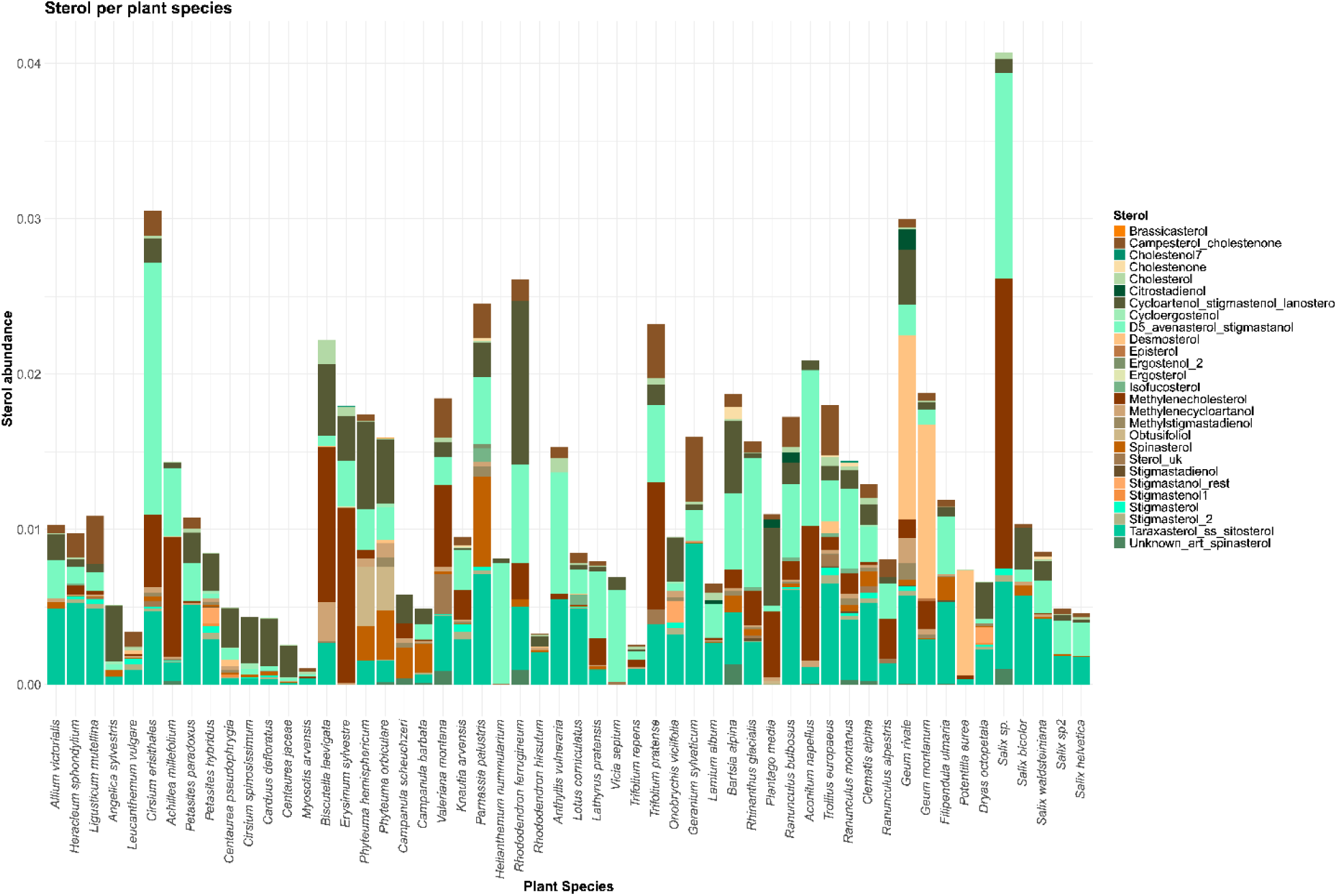
Composition of sterols in mg/g pollen of 54 plant species. Sterols are color-coded.

### Nutrient composition of collected and non-collected pollen

The FA composition of pollen from collected and non-collected plant species was marginally significantly different (F = 2.027, R^2^ = 0.027, df = 1, *p* = 0.092, 999 Permutations, Figure 6), particularly with regard to the combination of linolenic acid (ω-3, essential FAs, polyunsaturated) with oleic acid (monounsaturated, R^2^ = 0.54, *p* = 0.001), palmitic acid (R^2^ = 0.45, *p* = 0.001), stearic and octadecatrienoic acid (R^2^ = 0.37, *p* = 0.001) and linoleic (ω-6, essential FA, polyunsaturated, R^2^ = 0.34, *p* = 0.001). Collected pollen contained, on average, 21.3% less FA (69.8mg/g) than non-collected pollen, (91.1 mg/g). However, the difference was not significant (W = 616, *p* = 0.253, Figure 7A).

**Figure 6:**
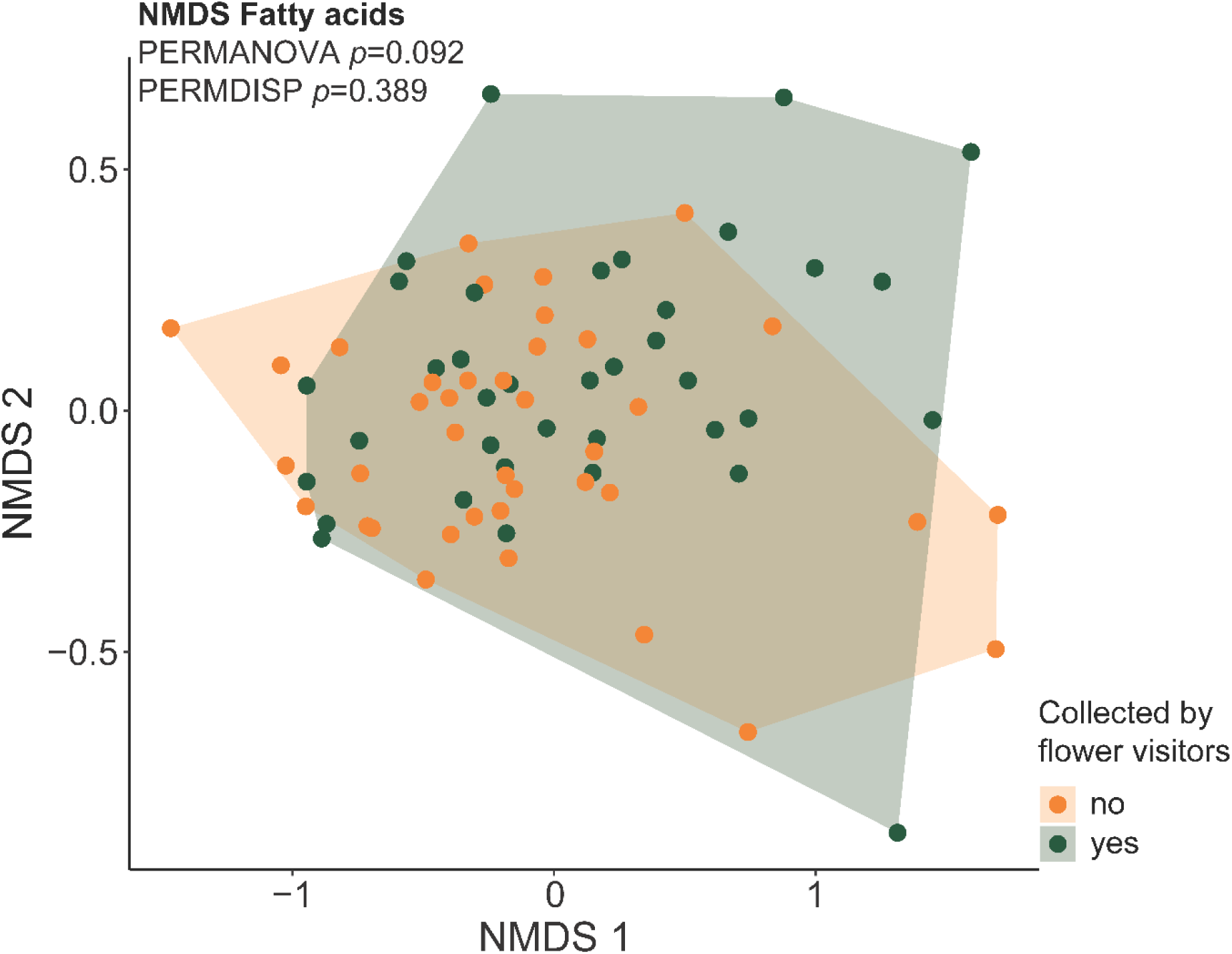
Composition of fatty acids in pollen collected by flower visitors (green) or not collected (orange), displayed by non-metric multi-dimensional scaling (NMDS).

**Figure 7:**
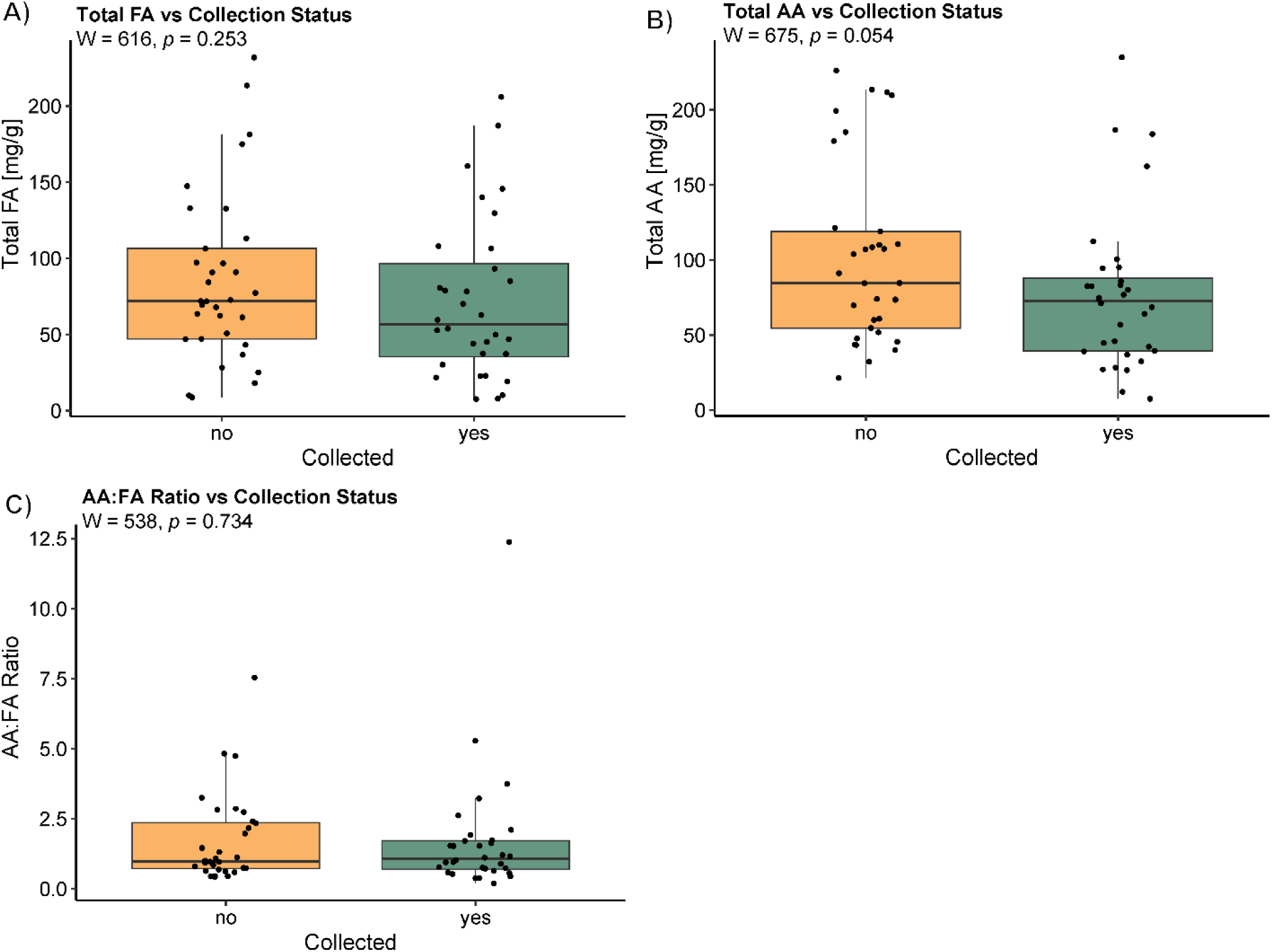
A) Total FA content of pollen, which was not collected by bumblebees and hoverflies (orange) and pollen collected by them (green). B) Total AA content of pollen, which was not collected by bumblebees and hoverflies (orange) and pollen collected by them (green).C) Pollen amino acid to fatty acid ratio of pollen not collected by bumblebees and hoverflies (orange) and pollen collected by them (green).

The AA composition was also marginally significantly different between collected and non-collected pollen (F = 2.972, R^2^ = 0.039, df = 1, *p* = 0.066, 999 Permutations, Figure 8). Collected pollen had, on average, 10.4% less AA (90.4 mg/g) than not collected pollen (100.9 mg/g; W = 675, *p* = 0.054, Figure 7B). Moreover, AA and FA were positively correlated in the analyzed pollen samples (R = 0.4, *p* = 0.001, Figure 9), while the P:L ratio of collected and not collected pollen did not differ (W = 538, *p* = 0.734, Figure 7C).

**Figure 8:**
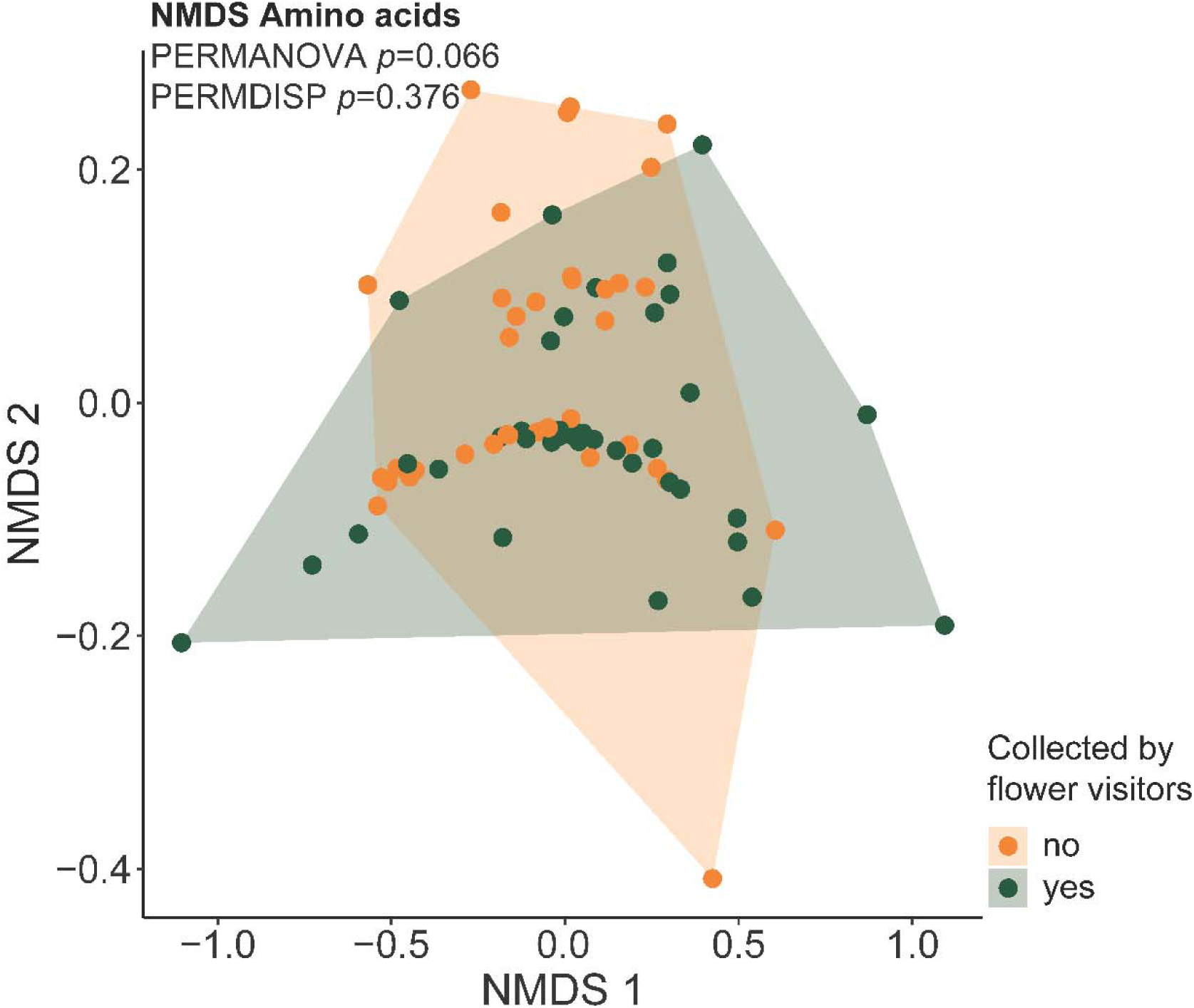
Composition of amino acids in pollen collected by flower visitors (green) or not collected (orange), displayed by Non-metric multi-dimensional scaling (NMDS).

**Figure 9:**
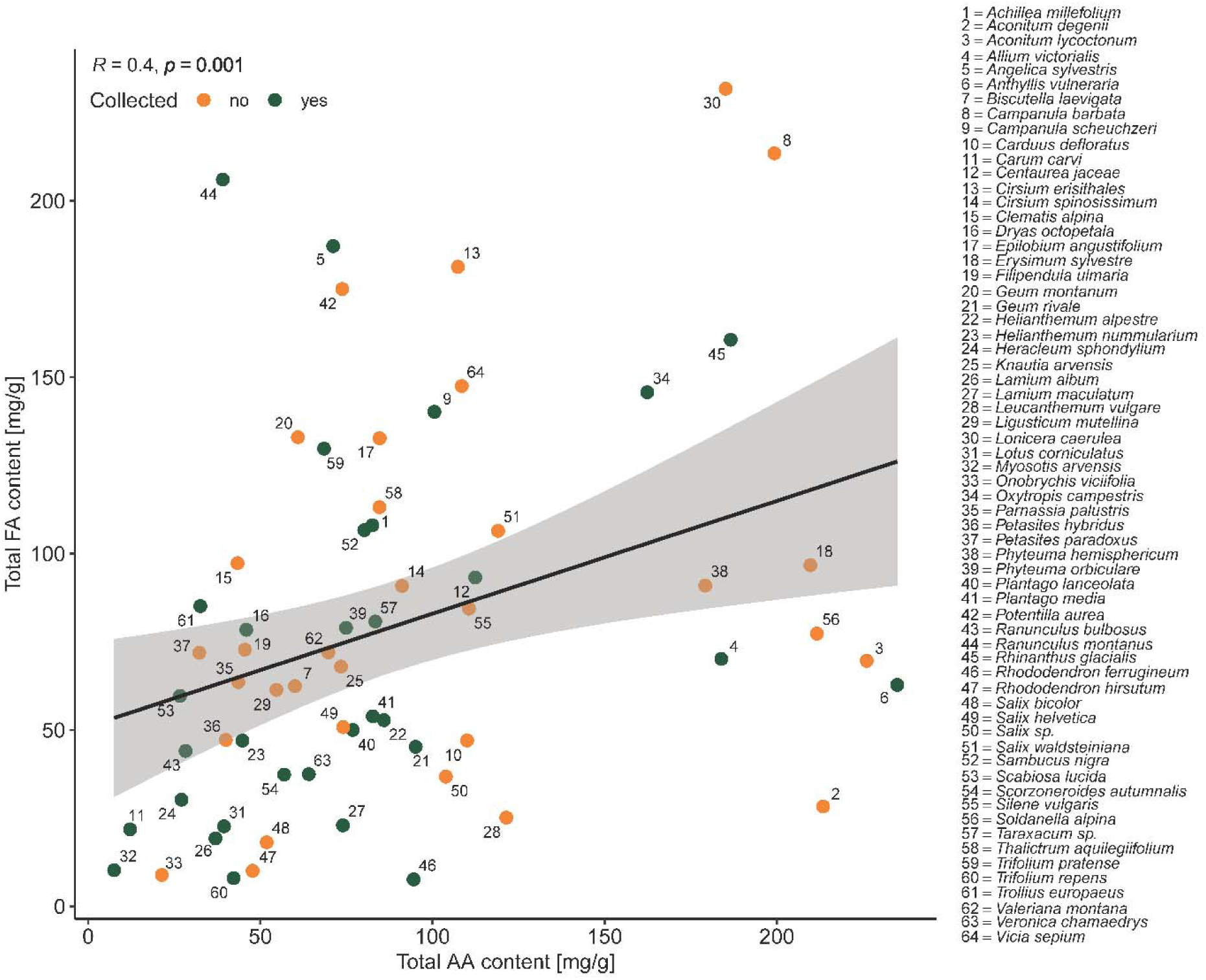
Correlations between total amino acid (AA) content and total fatty acid (FA) content. Green dots indicate collected pollen, orange dots non-collected pollen. The solid line shows the correlation between AA and FA content in pollen. Numbers represent pollen origins, as listed on the right.

Sterol composition and content were similar between collected and non-collected plant species (F = 0.498, R2 = 0.001, df = 1, *p* = 0.867, 999 Permutations, Figure 10). Pollen collected by bumblebees and hoverflies had an average sterol content of 12.7 µg/g, which was similar to pollen that was not collected (12.0 µg/g).

**Figure 10:**
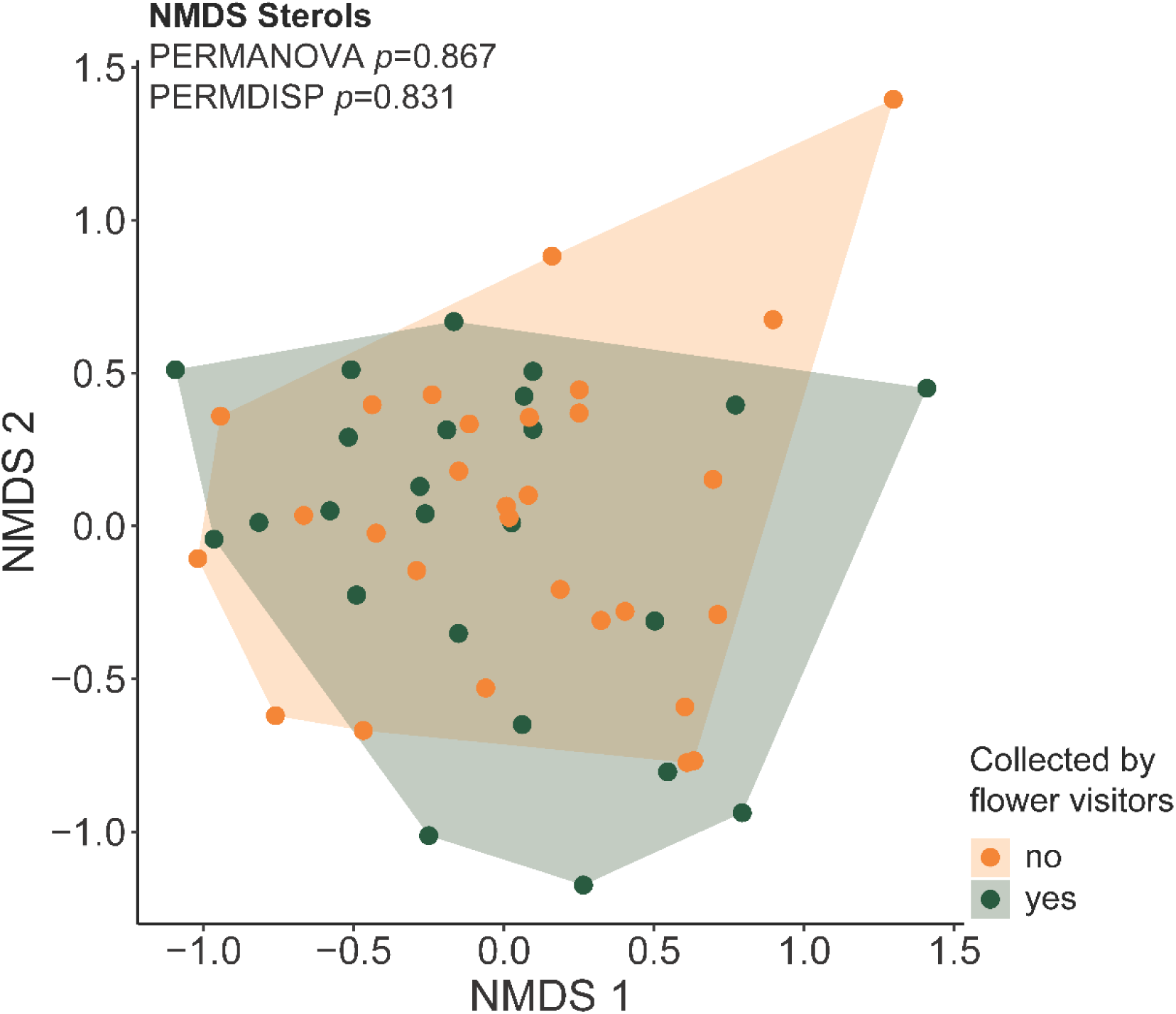
Composition of sterols in pollen collected by flower visitors (green) or not collected (orange), displayed by NMDS

### Phylogeny

Only one FA, arachidonic acid, showed a weak phylogenetic signal (K = 0.656, *p* = 0.001, N = 73; Figure 11), while no phylogenetic signal was detected for any other FA, nor for any of the AAs or sterols (Figure S4, Figure S5).

**Figure 11:**
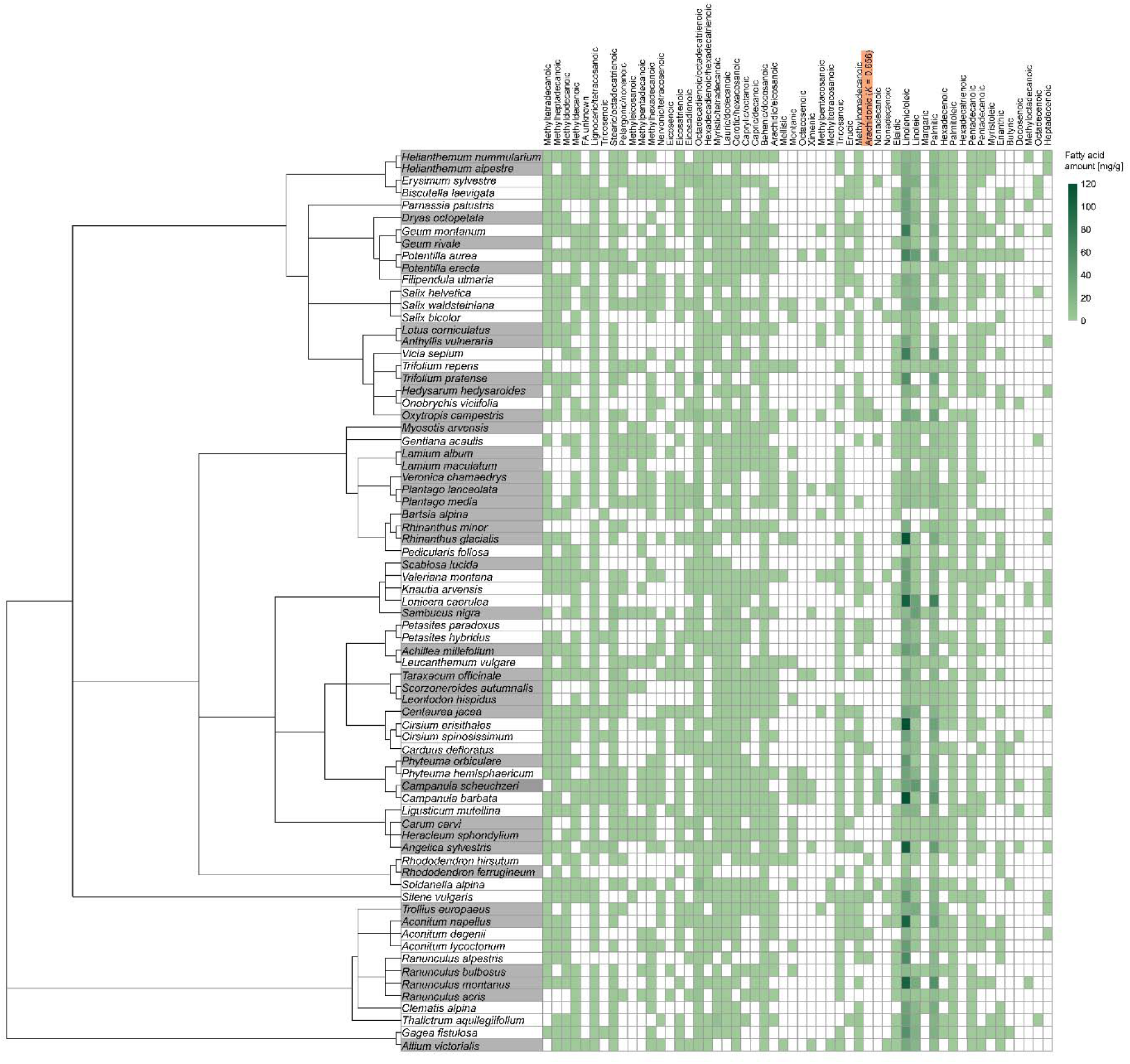
Pollen fatty acid profiles of plant species. Fatty acids are shown at the top with different concentrations depicted by different shades of green; the only fatty acid with a significant phylogenetic signal is marked in orange (Blomberg’s K). Phylogenetic relationships are given on the left. Plant species with a grey background were collected by pollinators; plant species with a white background were not collected.

## DISCUSSION

Although bees are known to exhibit nutrient-specific perception and regulation of pollen under laboratory conditions, nutrient-specific foraging behavior has hardly been observed in the field. By combining metabarcoding of pollen collected by alpine bumblebees and hoverflies with detailed analyses of pollen nutrient composition, we show that the two pollinator groups exploit a more similar range of floral resources than expected. Both groups collected pollen (respectively fed on pollen) with comparatively low FA and AA concentrations, suggesting that not only AA content alone, but in combination with FA content, acts as a common driver determining pollen collection in pollinators foraging in a natural system. In contrast, sterols —although physiologically essential— appear to play a minor role in shaping pollen-collection patterns.

We found that observed flower visit network and pollen-based metabarcoding-derived network differed significantly, indicating that flower visitors do not necessarily collect pollen from those plant species on which they are observed, and that they collect pollen from many additional species that couldn’t be observed in the field. Thus, observational data alone are insufficient to provide a reliable picture of the flowering plant species used by pollinators for pollen collection (Parreno et al., 2024). Our study also confirms previous recommendations on the use of both observational and pollen/metabarcoding-based data to obtain a complete picture of interactions (Bell et al., 2019; Pornon et al., 2016).

In the pollen-based network, almost all plants visited by bumblebees were also visited by hoverflies, but hoverflies collected pollen from a broader range of plant species. Hoverflies are likely constrained to open flowers with easily accessible nectaries and pollen sacs due to their short, dipping proboscis (Branquart & Hemptinne, 2000; Doyle et al., 2020; Klecka et al., 2018). However, we found them to collect pollen from almost any plant species in our dataset, including those also exploited by bumblebees. In contrast, in the network based on flower observations, some plant species were visited exclusively by bumblebees or by hoverflies. This incongruence may reflect differences in resource attractiveness: some plants may provide more appealing nectar, whereas others may be primarily attractive for their pollen (Leponiemi et al., 2023). Additionally, during nectar collection, pollinators may unintentionally pick up small amounts of pollen from the nectar-producing flowers (Westrich, 2018). Alternatively, methodological factors may explain the observed discrepancies: Observational sampling captures only a single behavioral event, whereas pollen metabarcoding integrates pollen collected across the insect’s entire foraging interval, or even up to several days in case of gut pollen samples (Borrero et al., 2026; Pornon et al., 2016).

Our initial hypothesis that the two focal groups of flower visitors exploit distinct floral plants was not supported. Instead, the results indicate a strong overlap in their nutritional niches, with hoverflies potentially utilizing a broader niche than bumblebees. Both primarily forage on plants with relatively low concentrations of FAs and AAs - a finding that partly aligns with previous laboratory studies showing that bumblebees and honeybees exhibit a high sensitivity for FAs (Ruedenauer et al., 2021; Ruedenauer et al., 2020; Ruedenauer et al., 2024; Schleifer et al., 2024): they not only perceive and discriminate among individual FAs, but also avoid pollen with elevated FA levels, even when this avoidance results in underconsumption of other essential nutrients and reduced survival (Ruedenauer et al., 2020; Ruedenauer et al., 2024; Schleifer et al., 2024). Despite the detrimental effect of comparatively high concentrations of FA on pollinators’ health (Ruedenauer et al., 2020; Schleifer et al., 2024), FAs are essential for most pollinators, including bees and hoverflies (Leonhardt et al., 2024). FAs play a pivotal role as an energy source, contribute to the integrity of membrane structure (Hulbert & Abbott, 2011) and cellular homeostasis, and are involved in cognitive processes (Arien et al., 2018; Arien et al., 2020; Arien et al., 2015). FAs also show antibiotic properties against several pathogens (Feldlaufer et al., 1993). However, excessive FA intake restrict absorption by midgut cells (Canavoso et al., 2001), potentially leading to cell membrane damage (Haddad et al., 2007). Furthermore, a high intake of ω-6 FAs (linoleic acid) interferes with the desaturation of ω-3 FAs (α-linolenic acid) (Patterson et al., 2012). The multiple roles and often contrasting effects of FAs may require their regulation within a tight concentration window (Simpson & Raubenheimer, 2012).

Notably, our results reached significance only for total AA content, but not for FA, even though pollen collected by pollinators contained on average 21.3 % less FA than pollen not collected by pollinators. However, total AA and total FA in pollen were clearly positively correlated across plant species, enabling pollinators to regulate both nutrients simultaneously. While previous results showed that the bumblebee *Bombus terrestris* and honeybees regulate FA content when foraging for pollen in laboratory experiments (Ruedenauer et al., 2020; Schleifer et al., 2024), it is unknown whether other bumblebee species and hoverflies also regulate FAs. For example, honeybees do perceive differences in AA concentrations in pollen, while *B. terrestris* workers do not, highlighting species-specific differences in nutrient perception – a prerequisite for regulation. However, if bumblebees and hoverflies in alpine ecosystems regulated pollen AA intake, it is counterintuitive that they collected pollen with comparatively low AA concentrations, because lower AA content are likely unfavorable for bees, which in general forage to increase protein intake and decrease FA intake (Kämper et al., 2016; Ruedenauer et al., 2019a; Ruedenauer et al., 2020; Ruedenauer et al., 2016; Ruedenauer et al., 2019b; Schleifer et al., 2024; Vaudo et al., 2016a; Vaudo et al., 2016b; Vaudo et al., 2020). AAs are essential for protein synthesis (Chapman, 1998), larval growth (De Groot, 1952), and as an energy source for the flight muscles (Micheu et al., 2000). However, previous findings show that bumblebees and honeybees largely tolerate fluctuations in AAs, as naturally observed in floral pollen, and do not regulate their intake (Ruedenauer et al., 2021; Ruedenauer et al., 2019; Ruedenauer et al., 2020).

Previous studies emphasized the importance of the protein-to-lipid (P:L) ratio for pollen consumers (Crone & Grozinger, 2021; Vaudo et al., 2016a). For example, Vaudo et al. (2016a) showed that pollen with high P:L ratios is generally favored by bumblebees up to a certain level for total FA concentrations. Given that, e.g., bumblebees and honeybees avoid high pollen fat concentrations (Ruedenauer et al., 2021; Ruedenauer et al., 2020; Schleifer et al., 2024), this previously observed preference for a high P:L ratio may be driven not by high protein levels but rather by low fat levels, as suggested by Ruedenauer et al. (2020). As AA concentrations correlated with FA concentrations in pollen in our study system, which was also found by previous studies (Roulston & Cane, 2000; Ruedenauer et al., 2019b; Vaudo et al., 2020), a preference for pollen with low fat content automatically resulted in low AA concentrations. However, the large variation in pollen FA content likely prevented statistical significance and indicates that pollen collection is affected by additional factors besides pollen FA and AA content. For example, other pollen compounds, e.g., plant secondary metabolites, minerals and their ratios (e.g., NA:K (Filipiak et al., 2023)), likely affect flower choices of pollinators. Moreover, differences in the floral density, abundance and general availability likely affect foraging choices (Fowler et al., 2016).

Beyond nutritional cues, other floral traits play also a crucial role in shaping pollinator foraging decisions. Visual cues such as flower color and contrast influence detectability and preferences, sometimes linked to sensory biases and learning abilities (Chittka & Menzel, 1992; Dyer et al., 2012). Floral morphology further constrains access to rewards, with corolla depth, symmetry, and mechanical complexity acting as filters that favour pollinators with matching body size or feeding structures (Fenster et al., 2004; Stang et al., 2006). In addition, floral scent provides long-distance signals that guide pollinators to rewarding resources and can reinforce constancy once profitable flowers are identified (Raguso, 2008; Wright & Schiestl, 2009). This indicates that, in natural environments, pollen foraging choices are not influenced solely by a single nutrient group, but rather by a combination of interacting factors.

Sterols were not at all linked to pollen collection in our study, although crucial for the structural integrity of the cell’s phospholipid bilayer (Clayton, 1964), as precursors for important steroid hormones, and for organismal growth and patterning (Clayton, 1964; Robbins et al., 1971; Svoboda et al., 1978). Additionally, most insects, including bees and hoverflies, cannot synthesize sterols de novo and need to obtain them via their food (Canavoso et al., 2001; Svoboda et al., 1978). This suggests that, although sterols are important, flower visitors do not prioritize their regulation (Nebauer et al., 2023), likely because these nutrients are provided in sufficient quantities in natural pollen and are not detrimental if consumed in comparatively high amounts. Regulation of many or even all nutrients is highly unlikely, as it would be extremely costly in terms of reception, perception, and behavioral adaptation (Bernays, 2001; Ruedenauer et al., 2023). It is thus plausible that organisms restrict nutrient regulation to those nutrients for which they show a narrower tolerance range (Simpson & Raubenheimer, 2012). This may also explain why animals follow a rule of compromise, when food is nutritionally imbalanced and animals cannot meet all nutrient targets simultaneously. In this case, they typically avoid overconsumption of the nutrient with the higher physiological cost, e.g., FA or toxic secondary compounds, even at the expense of underconsuming other nutrients (Raubenheimer & Simpson, 1999; Simpson & Raubenheimer, 2012; Simpson et al., 2004).

We had further expected that variation in pollen nutrients is shaped by flower visitation patterns rather than by plant phylogeny. And indeed, we did not find significant associations for any nutrient group. While previous studies found sterol and AA variation to be correlated with plant phylogeny (Mondal & Mandal, 2009; Zu et al., 2021), the lack of a respective correlation in our system might be due to strong environmental filtering in alpine ecosystems. Alpine plants experience extreme abiotic pressures, including low temperatures, intense UV radiation, recurrent frost events, and very short growing seasons, which exert strong selection on functional traits across unrelated lineages (Körner, 1999). Such harsh conditions frequently drive ecological convergence, whereby species evolve similar reproductive, physiological, or biochemical traits despite distant phylogenetic relationships (Losos, 2011). Given that pollen biochemistry is sensitive to environmental stress, particularly temperature and desiccation that ia known to affect AA concentrations in pollen (Descamps et al., 2021), it is plausible that a similar selective pressure across alpine habitats promotes convergence in pollen nutrient profiles across lineages.

## CONCLUSION

Alpine bumblebees and hoverflies share a similar taxonomic and nutritional niche, both favoring pollen from plants with comparatively low concentrations of AAs and FAs. Our findings suggest that nutrient concentrations in food contribute to but are not the only driver of plant-pollinator interactions in natural ecosystems.

## AUTHOR CONTRIBUTION

SDL, JS, FAR, JN, and MCS conceived the ideas and designed the methodology; MSC and LC collected the data; MCS and AK analyzed samples and data; MCS led the writing of the manuscript. All authors contributed critically to the drafts and gave final approval for publication.

## ACKNOWLEDGEMENT

We thank Stefanie Siebler for help with the AA analysis and Anika Schopf for the metabarcoding analysis. We also want to thank the Großglockner Hochalpenstraßen AG and the Haus der Natur in Salzburg for using the Eberhardt Stübner Forschungsstation. We finally thank the Deutsche Forschungsgemeinschaft (DFG project: LE 2750/5-2 to SDL and SP1380/1-2 to JS) for financial support. This research was conducted with the approval of the Bezirkshauptmannschaft Spittal an der Drau (approval number SP3-NS_4157/2023).

## STATEMENT OF INCLUSION

Our research was conducted entirely by researchers based in Germany and Austria and did not involve data collection in other regions. The authorship, therefore, reflects the regional context of the research, and relevant literature was considered where appropriate.

## DATA AVAILABILITY

All datasets generated and analyzed during this study will be publicly available upon publication.

## TRANSPARENCY DISCLOSURE

AI (Grammarly, Microsoft Copilot, and ChatGPT) was used to improve statistical analyses, coding, and language, but not for text generation. The authors made all intellectual contributions, interpretations, and conclusions.

