## Supplementary Material for "No need for fat: Nutritional preferences affect pollen foraging networks in alpine pollinator communities"

**S1 Analyis of pollen AAs:**

Free and protein-bound AA analysis of the pollen samples by ion exchange chromatography (IEC, Amino Acid Analyzer LC3000, Eppendorf-Biotronic, Hamburg, Germany) was performed following Leonhardt and Blüthgen (2011). To analyze the water-soluble AAs, 5–10 mg of pollen (dry weight) was mixed with 100–200 µl of bidistilled H₂O and placed in an ultrasonic bath for 30 minutes, followed by a 1-hour extraction in the refrigerator. After 5–10 minutes of centrifuging, the supernatant was transferred into a fresh microcentrifuge tube, and the sediment was saved for later protein-bound AA analysis. The supernatant was boiled at 100°C for 2 minutes in a heating block and then cooled on ice to room temperature. After centrifuging for 5 minutes, 50 µl of the supernatant was mixed with 10 µl of 12.5% sulphosalicylic acid (Sigma-Aldrich, St. Louis, MO, USA) for precipitation and extracted in the refrigerator for 30 minutes. The solution was then mixed, centrifuged again for 10 minutes, and 50 µl of the supernatant was combined with 50 µl of sample dilution buffer (lithium loading buffer, Laborservice Onken, Gründau, Germany) and applied to a membrane filter (Vecta Spin). After centrifuging for 5 minutes, the sample was analyzed by IEC. To analyze the protein-bound AAs, the sediment extracted in the first step was mixed with 200 µl of 6 N HCl and boiled for 4 hours at 100°C in a heating block, followed by cooling on ice to room temperature. After centrifuging for 10 minutes, the supernatant was transferred into a new microcentrifuge tube and evaporated completely at 100°C in the heating block. Then, 200 µl of bidistilled H₂O was added, and the mixture was boiled again at 100°C for 30–45 minutes. An additional 200 µl of bidistilled H₂O was then added. After cooling on ice to room temperature, the mixture was centrifuged for 10 minutes. Of this supernatant, 100 µl was mixed with 20 µl of 12.5% sulphosalicylic acid and extracted in the refrigerator for 30 minutes. The solution was mixed and centrifuged for 10 minutes. In a fresh microcentrifuge tube, 100 µl of the supernatant was mixed with 100 µl of sample dilution buffer and transferred onto a membrane filter. After centrifuging for 5 minutes, the solution was further diluted by mixing 20 µl of the sample solution with 80 µl of sample dilution buffer and was then ready for IEC analysis.

**S2 Analyis of pollen fatty acid:**

Pollen FA content was analyzed based on Villagómez et al. (2023). Therefore, 0.5–1.00 mg of pollen for each sample was weighed into a 2 mL vial, to which 7 µL of FA standard (nonadecanoic acid in methanol, 0.2 mg/mL, Sigma-Aldrich, St. Louis, MO, USA; absent in pollen and other plant tissues) and 0.5 mL of a chloroform-methanol mixture (2:1, Sigma-Aldrich, St. Louis, MO, USA) were added. A glass rod was used to disrupt the pollen exines until the pollen grains were nearly undetectable in the solvent. The vial was then filled with the chloroform-methanol mixture to achieve a final volume of 1.5 mL and transferred into a larger glass vial (minimum volume of 3 mL). An additional 1.5 mL of the chloroform-methanol mixture was used to rinse the vial, and this rinse was combined with the previously transferred solution in the larger glass vial. Subsequently, the large glass vials were placed in a thermocycler and shaken for 24 hours at room temperature at 300 rpm. The following day, once the incubation period was complete and the pollen had settled, excess solvent was evaporated under controlled nitrogen (N2) airflow until only a tiny amount of liquid remained. The supernatant was transferred into a 300 µL inlet within the vial and completely evaporated under N2 airflow. The samples were stored at -20°C until further analysis. Before gas chromatography-mass spectrometry (GC-MS) analysis, 20 µL of Trimethylsulfonium hydroxide (TMSH, Macherey Nagel, Düren, Germany) was added as a FA derivatization agent to convert the FAs into FA methyl esters (FAMEs). This mixture was vortexed and then analyzed using the GC-MS (GC: 6890N, MSD: 5975, Agilent Technologies, Santa Clara, United States). To identify the FAs, their mass spectra, retention times, and retention indices were analyzed and compared to the area of the FA standard (nonadecanoic acid). The total FA content was calculated by summing all FAs. A control (blank sample run without pollen) was conducted to assess any potential FA contamination that may have occurred during the process. The calculated concentration was subtracted from the sample profiles if contamination was identified.

**S3 Analyis of pollen sterols:**

The sterol content of pollen was analyzed according to the method described by Vanderplanck et al. (2011). Approximately 15 to 20 mg of pollen was weighed into a round-bottom flask and mixed with 2.5 mL of 2M methanolic KOH (Sigma-Aldrich, St.Louis, MO, USA). The mixture underwent saponification for 1 hour in a water bath maintained at 80°C, equipped with a cooler set at 15°C. After saponification, the mixture was allowed to cool for about 10 minutes, and then 4 mL of a betulin standard (Sigma-Aldrich, St.Louis, MO, USA, equivalent to 0.4 mg betulin) was added. This solution was subsequently diluted with 10 mL of deionized water. The mixture was then transferred to a separatory funnel and shaken. The saponification apparatus was rinsed with 10 mL of diethyl ether (Sigma-Aldrich, St.Louis, MO, USA), which was added to the separatory funnel to extract the sterols. After allowing the phases to separate, the upper ether phase was collected in a new round-bottom flask. This extraction process was repeated two more times. The collected ether solution was washed three times with deionized water in the separatory funnel, then dried over anhydrous Na₂SO₄ (Sigma-Aldrich, St.Louis, MO, USA) in the round-bottom flask. Care was taken to transfer the ether to a new round-bottom flask without carrying over the Na₂SO₄. Following the complete evaporation of the diethyl ether, the residue was dissolved in 1-2 mL of hexane (Sigma-Aldrich, Burlington, MA, USA) and transferred to a 1.5 mL glass vial. The hexane was then entirely evaporated under a nitrogen (N₂) stream. This sample was stored at -20°C until further analysis. Before gas chromatography/mass spectrometry (GC-MS) analysis, the residue was mixed with 100 µL of anhydrous pyridine (Sigma-Aldrich, Burlington, MA, USA) and 100 µL of BSTFA (Sigma-Aldrich, St.Louis, MO, USA) in a thermomixer at 90°C and 1400 rpm for 30 minutes. After the reaction, the reagent evaporated, and the residue was dissolved in 100 µL of hexane. This final mixture was vortexed and subsequently analyzed using GC-MS (GC: 6890N, MSD: 5975, Agilent Technologies, Santa Clara, United States).

Tables and Figures

Table S 1: Plant species included in this study. For each species, the presence of bumblebees or hoverflies as floral visitors is indicated with “yes”; “no” indicates that the insect was not observed on this plant species. “NA” means that the plant species was not observed. If pollen of a given plant species was detected in the pooled pollen loads of bumblebees or hoverflies, this is likewise marked with “yes”; if no pollen was found, “no” is shown. Nutrient content (FAs, Aas, and sterols) is reported in µg/mg. “NA” indicates that pollen quantity was insufficient for analysis.

| **Plant species** | | **Observation** | | **Barcoding** | | **Nutrients [µg/mg]** | | |
| --- | --- | --- | --- | --- | --- | --- | --- | --- |
| **Species** | **Family** | **Bees** | **Flies** | **Bees** | **Flies** | **FAs** | **Sterols** | **AAs** |
| *Achillea millefolium* | Asteraceae | no | no | no | **yes** | 108.04 | 0.0144 | 82.54 |
| *Aconitum degenii* | Ranunculaceae | **yes** | **yes** | no | no | 28.31 | NA | 213.44 |
| *Aconitum lycoctonum* | Ranunculaceae | no | no | no | no | 69.58 | NA | 226.10 |
| *Aconitum napellus* | Ranunculaceae | **yes** | no | **yes** | **yes** | 156.21 | 0.0209 | 283.13 |
| *Aconitum variegatum* | Ranunculaceae | NA | NA | **yes** | **yes** | NA | NA | NA |
| *Aegopodium podagraria* | Apiaceae | NA | NA | no | **yes** | NA | NA | NA |
| *Allium victorialis* | Amaryllidaceae | **yes** | **yes** | **yes** | **yes** | 70.14 | 0.0103 | 183.88 |
| *Angelica sylvestris* | Apiaceae | no | no | no | **yes** | 187.14 | 0.0051 | 71.16 |
| *Anthoxanthum alpinum* | Poaceae | NA | NA | no | **yes** | NA | NA | NA |
| *Anthyllis vulneraria* | Fabaceae | **yes** | no | **yes** | **yes** | 62.76 | 0.0153 | 234.97 |
| *Aster alpinus* | Asteraceae | NA | NA | no | **yes** | NA | NA | NA |
| *Aster* sp. | Asteraceae | NA | NA | no | **yes** | NA | NA | NA |
| *Asteraceae* sp. | Asteraceae | NA | NA | no | **yes** | NA | NA | NA |
| *Bartsia alpina* | Orobanchaceae | no | no | **yes** | **yes** | 0.47 | 0.0187 | 152.98 |
| *Biscutella laevigata* | Brassicaceae | no | no | no | no | 62.43 | 0.0222 | 60.06 |
| *Bromus* sp. | Poaceae | NA | NA | no | **yes** | NA | NA | NA |
| *Calluna vulgaris* | Ericaceae | **yes** | no | **yes** | **yes** | NA | NA | 65.11 |
| *Campanula barbata* | Campanulacea | **yes** | no | no | no | 213.48 | 0.0048 | 199.27 |
| *Campanula scheuchzeri* | Campanulacea | **yes** | no | **yes** | **yes** | 140.18 | 0.0057 | 100.56 |
| *Carduus crispus* | Asteraceae | NA | NA | **yes** | **yes** | NA | NA | NA |
| *Carduus defloratus* | Asteraceae | no | no | no | no | 47.00 | 0.0043 | 110.06 |
| *Carum carvi* | Apiaceae | NA | NA | no | **yes** | 21.81 | NA | 12.19 |
| *Centaurea jaceae* | Asteraceae | no | no | **yes** | **yes** | 93.23 | 0.0025 | 112.42 |
| *C. pseudophrygia* | Asteraceae | **yes** | **yes** | no | no | 412.07 | 0.0049 | 90.75 |
| *Cirsium erisithales* | Asteraceae | no | no | no | no | 181.27 | 0.0305 | 107.38 |
| *Cirsium spinosissimum* | Asteraceae | no | no | no | no | 90.84 | 0.0044 | 91.14 |
| *Clematis alpina* | Ranunculaceae | no | no | no | no | 97.28 | 0.0129 | 43.40 |
| *Crepis aurea* | Asteraceae | NA | NA | no | **yes** | NA | NA | NA |
| *Crepis biennis* | Asteraceae | NA | NA | no | **yes** | NA | NA | 64.08 |
| *Crepis chondrilloides* | Asteraceae | NA | NA | no | **yes** | NA | NA | NA |
| *Doronicum glaciale* | Asteraceae | NA | NA | no | **yes** | NA | NA | NA |
| *Dryas octopetala* | Rosaceae | no | **yes** | no | **yes** | 78.40 | 0.0066 | 45.92 |
| *Epilobium angustifolium* | Onagraceae | no | no | no | no | 132.69 | NA | 84.66 |
| *Erysimum sylvestre* | Brassicaceae | no | no | no | no | 96.70 | 0.0179 | 209.74 |
| *Euphrasia officinalis* | Orobanchaceae | no | **yes** | no | no | NA | NA | 67.95 |
| *Euphrasia rostkoviana* | Orobanchaceae | NA | NA | no | **yes** | NA | NA | NA |
| *Filipendula ulmaria* | Rosaceae | no | no | no | no | 72.74 | 0.0119 | 45.53 |
| *Gagea fistulosa* | Liliaceae | no | no | no | no | 117.06 | NA | NA |
| *Gentiana acaulis* | Gentianaceae | no | no | no | no | 35.88 | NA | NA |
| *Gentianella germanica* | Gentianaceae | NA | NA | no | **yes** | NA | NA | NA |
| *Geranium sylvaticum* | Geraniaceae | **yes** | **yes** | **yes** | **yes** | NA | 0.0159 | 40.33 |
| *Geum montanum* | Rosaceae | no | **yes** | no | no | 132.99 | 0.0188 | 60.91 |
| *Geum rivale* | Rosaceae | **yes** | **yes** | **yes** | **yes** | 45.21 | 0.0299 | 95.13 |
| *Geum urbanum* | Rosaceae | NA | NA | **yes** | **yes** | NA | NA | 52.85 |
| *Hedysarum hedysaroides* | Fabaceae | yes | no | **yes** | **yes** | 42.03 | NA | NA |
| *Helianthemum alpestre* | Cistaceae | no | no | **yes** | **yes** | 52.77 | NA | 85.90 |
| *Helianthemum nummularium* | Cistaceae | no | no | **yes** | **yes** | 46.97 | 0.0081 | 44.80 |
| *Helianthemum* sp. | Cistaceae | NA | NA | **yes** | **yes** | NA | NA | NA |
| *Heracleum sphondylium* | Apiaceae | NA | NA | no | **yes** | 30.24 | 0.0097 | 27.10 |
| *Hieracium* sp. | Asteraceae | NA | NA | no | **yes** | NA | NA | NA |
| *Homogyne alpina* | Asteraceae | NA | NA | no | **yes** | NA | NA | NA |
| *Hypericum maculatum* | Hypericaceae | NA | NA | **yes** | **yes** | NA | NA | NA |
| *Knautia arvensis* | Caprifoliaceae | no | no | no | no | 67.96 | 0.0095 | 73.43 |
| *Lamium album* | Lamiaceae | NA | NA | **yes** | **yes** | 19.24 | 0.0065 | 36.98 |
| *Lamium maculatum* | Lamiaceae | NA | NA | **yes** | **yes** | 22.94 | NA | 73.98 |
| *Lathyrus pratensis* | Fabaceae | no | no | no | no | NA | 0.0079 | 69.87 |
| *Leontodon hispidus* | Asteraceae | NA | NA | **yes** | **yes** | 30.23 | NA | NA |
| *Leontodon* sp. | Asteraceae | NA | NA | no | **yes** | NA | NA | NA |
| *Leucanthemum vulgare* | Asteraceae | NA | NA | no | no | 25.15 | 0.0034 | 121.42 |
| *Ligusticum mutellina* | Apiaceae | no | **yes** | no | no | 61.33 | 0.0108 | 54.68 |
| *Lonicera caerulea* | Caprifoliaceae | no | **yes** | no | no | 231.81 | NA | 185.14 |
| *Lotus corniculatus* | Fabaceae | **yes** | no | **yes** | **yes** | 22.68 | 0.0085 | 39.46 |
| *Meum athamanticum* | Apiaceae | NA | NA | no | **yes** | NA | NA | NA |
| *Myosotis alpestris* | Boraginaceae | NA | NA | **yes** | **yes** | NA | NA | NA |
| *Myosotis arvensis* | Boraginaceae | NA | NA | no | **yes** | 10.25 | 0.0011 | 7.55 |
| *Onobrychis viciifolia* | Fabaceae | no | no | no | no | 8.88 | 0.0095 | 21.41 |
| *Oxytropis campestris* | Fabaceae | **yes** | no | **yes** | **yes** | 145.70 | 0 | 162.35 |
| *Parnassia palustris* | Celastraceae | no | **yes** | no | no | 63.56 | 0.0245 | 43.62 |
| *Pedicularis elongata* | Orobanchaceae | NA | NA | **yes** | no | NA | NA | NA |
| *Pedicularis foliosa* | Orobanchaceae | **yes** | no | no | no | 7.38 | NA | NA |
| *Pedicularis* sp. | Orobanchaceae | NA | NA | **yes** | **yes** | NA | NA | NA |
| *Petasites hybridus* | Asteraceae | no | no | no | no | 47.16 | 0.0085 | 39.98 |
| *Petasites paradoxus* | Asteraceae | no | no | no | no | 71.89 | 0.0108 | 32.28 |
| *Phyteuma hemisphericum* | Campanulacea | no | no | no | no | 90.92 | 0.0174 | 179.21 |
| *Phyteuma orbiculare* | Campanulacea | **yes** | **yes** | **yes** | **yes** | 78.94 | 0.0158 | 74.89 |
| *Pilosella peleteriana* | Asteraceae | NA | NA | no | **yes** | NA | NA | NA |
| *Pimpinella major* | Apiaceae | NA | NA | no | **yes** | NA | NA | NA |
| *Plantago lanceolata* | Plantaginaceae | NA | NA | no | **yes** | 50.00 | NA | 76.82 |
| *Plantago media* | Plantaginaceae | NA | NA | no | **yes** | 53.90 | 0.0109 | 82.66 |
| *Poa* sp. | Poaceae | NA | NA | no | **yes** | NA | NA | NA |
| *Potentilla aurea* | Rosaceae | yes | yes | no | no | 175.00 | 0.0074 | 73.78 |
| *Potentilla erecta* | Rosaceae | NA | NA | no | **yes** | 12.72 | NA | NA |
| *Pulsatilla alpina* | Ranunculaceae | no | no | no | no | 67.60 | NA | NA |
| *Ranunculus acris* | Ranunculaceae | NA | NA | no | **yes** | 37.26 | NA | 475.30 |
| *Ranunculus alpestris* | Ranunculaceae | no | no | no | no | 82.82 | 0.0081 | NA |
| *Ranunculus bulbosus* | Ranunculaceae | NA | NA | no | **yes** | 44.04 | 0.0172 | 28.30 |
| *Ranunculus glacialis* | Ranunculaceae | NA | NA | no | **yes** | NA | NA | NA |
| *Ranunculus montanus* | Ranunculaceae | no | no | no | **yes** | 206.02 | 0.0144 | 39.10 |
| *Ranunculus polyanthemos* | Ranunculaceae | NA | NA | no | **yes** | NA | NA | NA |
| *Ranunculus spc* | Ranunculaceae | NA | NA | no | **yes** | NA | NA | NA |
| *Rhinanthus glacialis* | Orobanchaceae | **yes** | **yes** | no | no | 160.62 | 0.0156 | 186.61 |
| *Rhinanthus minor* | Orobanchaceae | NA | NA | **yes** | **yes** | 43.29 | NA | 107.11 |
| *Rhododendron ferrugineum* | Ericaceae | **yes** | **yes** | **yes** | **yes** | 7.63 | 0.0260 | 94.50 |
| *Rhododendron hirsutum* | Ericaceae | **yes** | **yes** | no | no | 10.07 | 0.0033 | 47.76 |
| *Rubus idaeus* | Rosaceae | NA | NA | no | **yes** | NA | NA | NA |
| *Salix bicolor* | Salicaceae | no | no | no | no | 18.12 | 0.0103 | 51.86 |
| *Salix hastata* | Salicaceae | NA | NA | no | no | NA | NA | 138.23 |
| *Salix helvetica* | Salicaceae | no | no | no | no | 50.80 | 0.0046 | 74.07 |
| *Salix herbacea* | Salicaceae | NA | NA | **yes** | **yes** | NA | NA | NA |
| *Salix* sp. | Salicaceae | **yes** | no | no | no | 36.77 | 0.0407 | 103.89 |
| *Salix* sp.2 | Salicaceae | **yes** | no | no | no | NA | 0.0049 | NA |
| *Salix waldsteiniana* | Salicaceae | no | no | no | no | 106.42 | 0.0086 | 119.05 |
| *Sambucus nigra* | Adoxaceae | NA | NA | no | **yes** | 106.61 | NA | 80.17 |
| *Saxifraga aizoides* | Saxifragecae | NA | NA | no | **yes** | NA | NA | NA |
| *Scabiosa lucida* | Caprifoliaceae | no | **yes** | **yes** | **yes** | 59.65 | NA | 26.67 |
| *Scorzoneroides autumnalis* | Asteraceae | NA | NA | **yes** | **yes** | 37.33 | NA | 56.92 |
| *Senecio nemorensis* | Asteraceae | NA | NA | no | **yes** | NA | NA | NA |
| *Senecio* sp. | Asteraceae | NA | NA | no | **yes** | NA | NA | NA |
| *Silene vulgaris* | Caryophyllaceae | no | **yes** | no | no | 84.38 | NA | 110.59 |
| *Soldanella alpina* | Primulaceae | no | no | no | no | 77.31 | NA | 211.65 |
| *Taraxacum sp.* | Asteraceae | no | **yes** | no | **yes** | 80.73 | NA | 83.38 |
| *Thalictrum aquilegiifolium* | Ranunculaceae | no | no | no | no | 113.15 | NA | 84.59 |
| *Thymus praecox* | Lamiaceae | NA | NA | **yes** | **yes** | NA | NA | NA |
| *Trifolium pratense* | Fabaceae | **yes** | **yes** | **yes** | **yes** | 129.73 | 0.0232 | 68.53 |
| *Trifolium repens* | Fabaceae | NA | NA | **yes** | no | 7.98 | 0.0027 | 42.23 |
| *Triticum turgidum* | Poaceae | NA | NA | no | **yes** | NA | NA | NA |
| *Trollius europaeus* | Ranunculaceae | **yes** | no | **yes** | **yes** | 85.09 | 0.0180 | 32.61 |
| *Vaccinium myrtillus* | Ericaceae | NA | NA | **yes** | **yes** | NA | NA | NA |
| *Vaccinium* sp. | Ericaceae | NA | NA | **yes** | no | NA | NA | NA |
| *Vaccinium uliginosum* | Ericaceae | NA | NA | **yes** | **yes** | NA | NA | NA |
| *Valeriana montana* | Caprifoliaceae | no | **yes** | no | no | 71.99 | 0.0185 | 69.76 |
| *Veronica chamaedrys* | Plantaginaceae | NA | NA | no | **yes** | 37.46 | NA | 64.09 |
| *Veronica serpyllifolia* | Plantaginaceae | NA | NA | no | **yes** | NA | NA | NA |
| *Vicia sepium* | Fabaceae | no | no | no | no | 147.47 | 0.0069 | 108.51 |

Relative Nutrient Compositions:

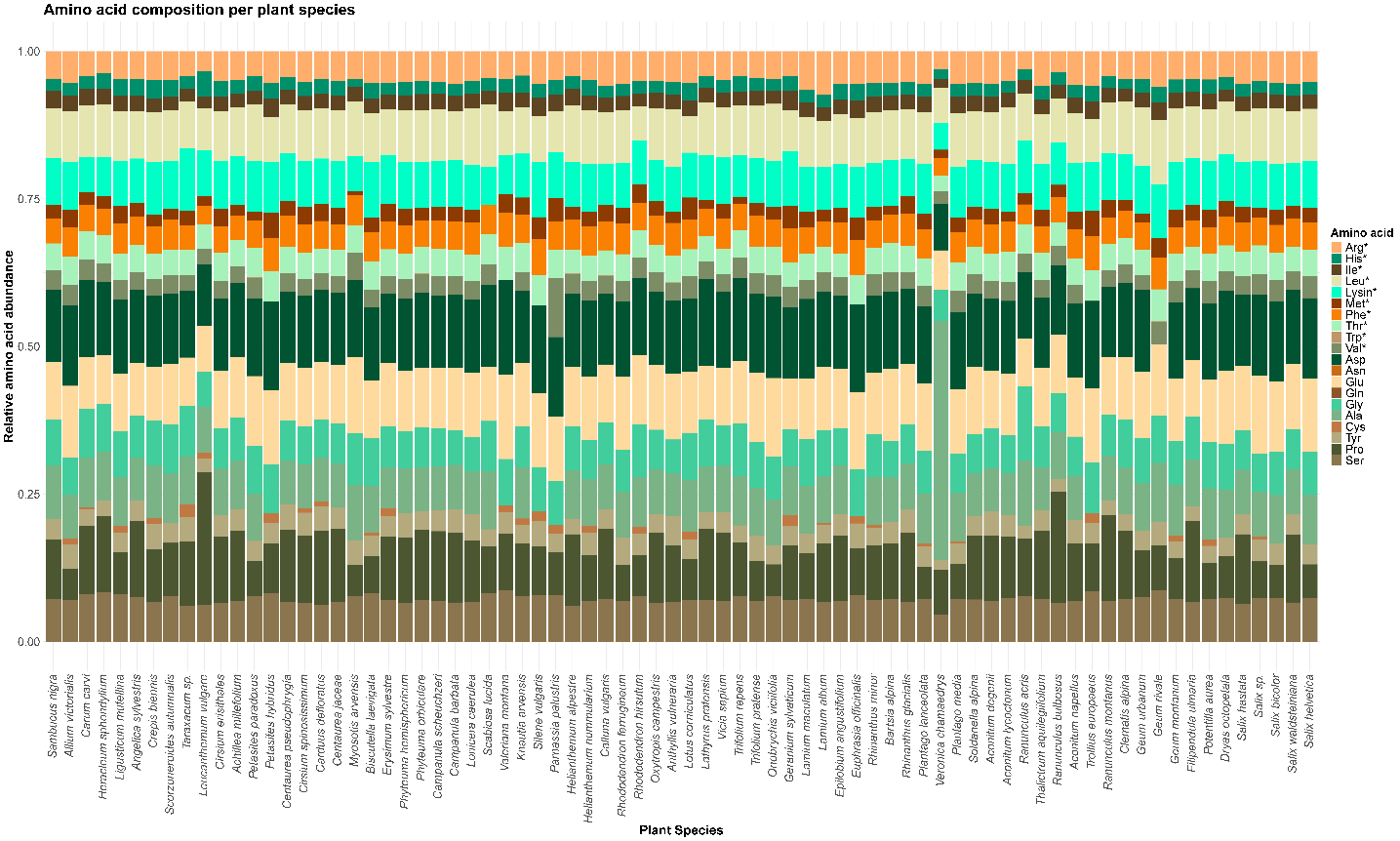

Figure S1: Relative composition of AAs in pollen of 75 plant species. AAs are color-coded. An asterisk after the name indicates essential AAs (EAA).

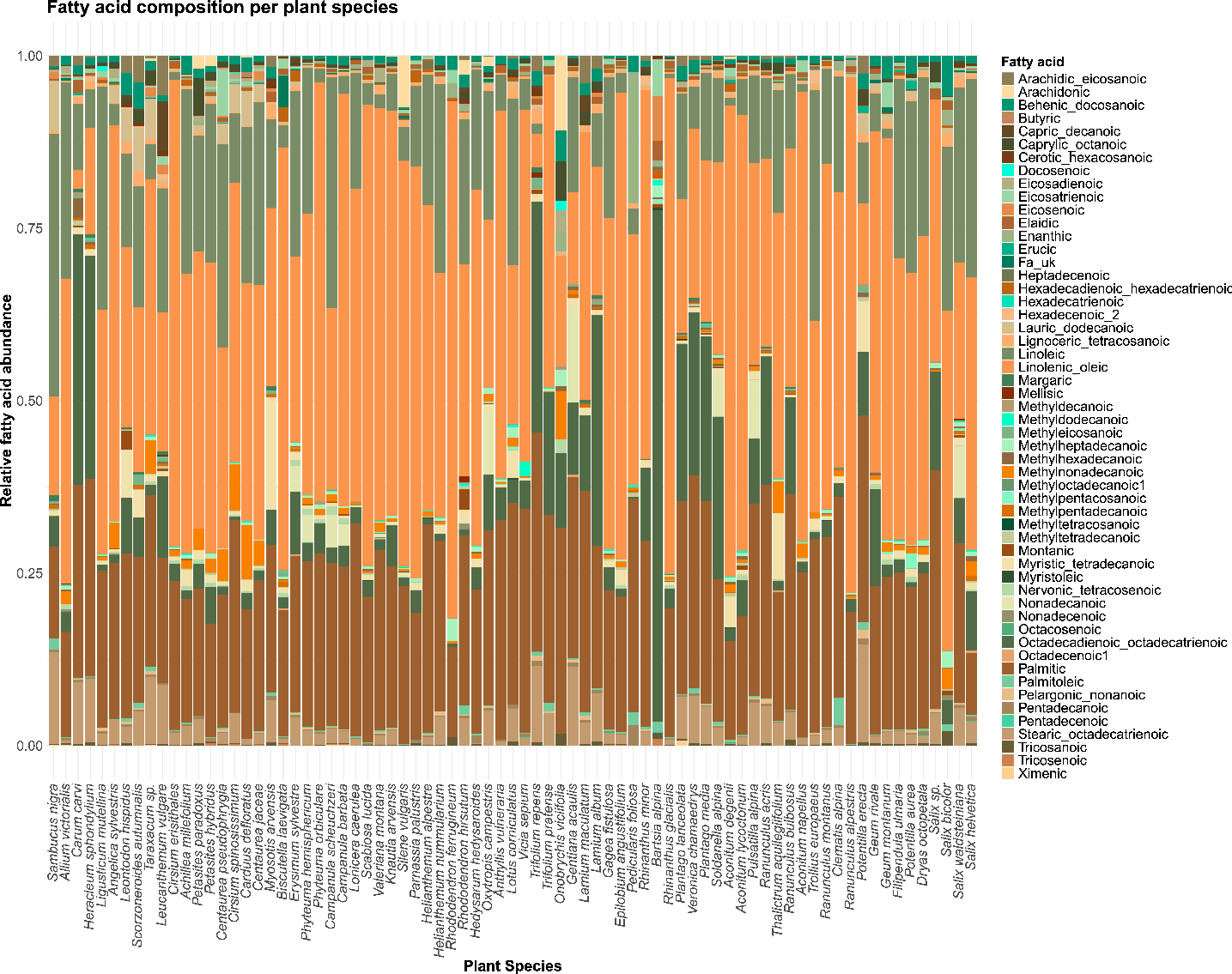

Figure S2: Relative composition of FAs in mg/g pollen of 77 plant species. FAs are color-coded. Linoleic and linolenic acids are considered essential FAs (EFAs).

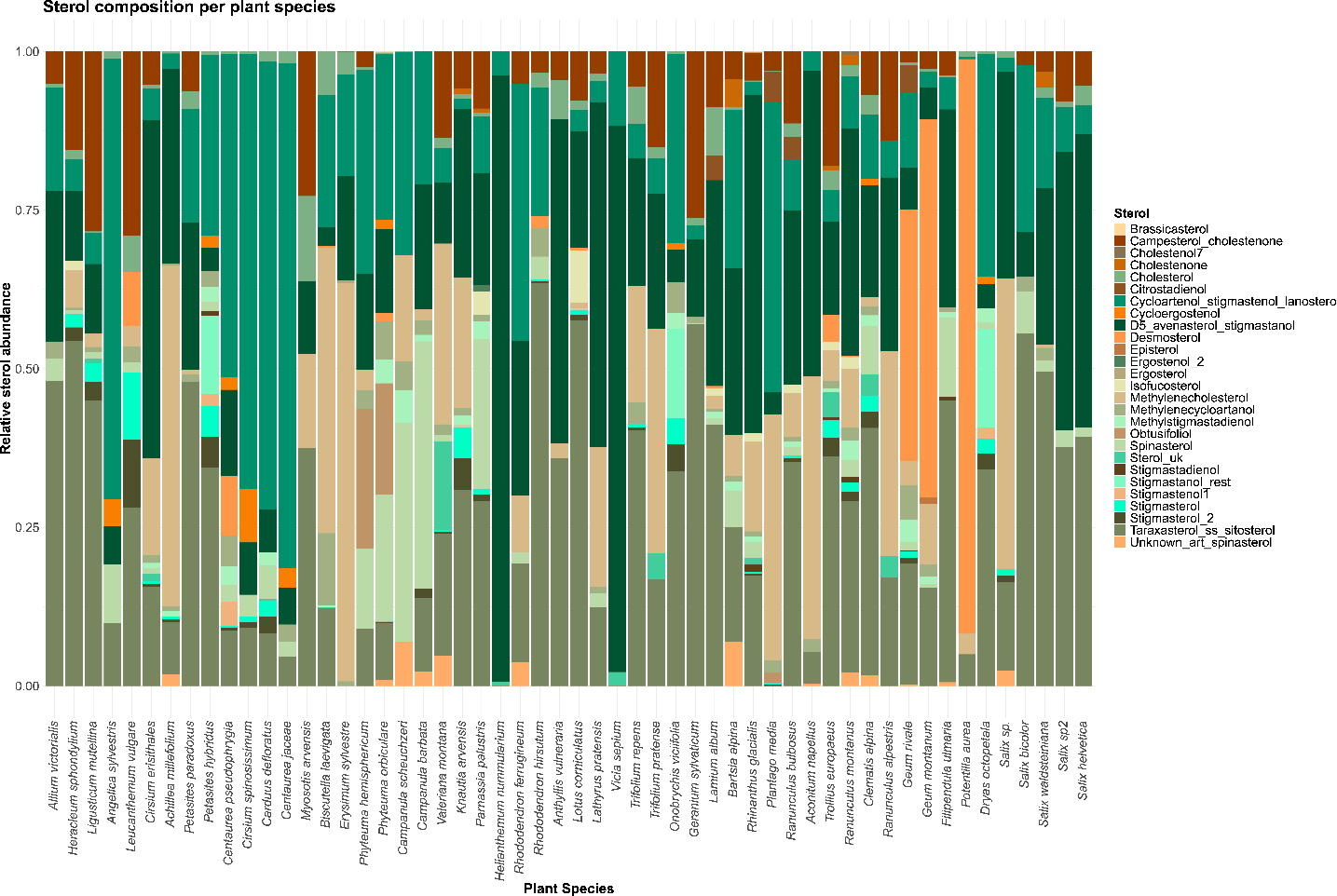

Figure S3: Relative composition of sterols in mg/g pollen of 54 plant species. Sterols are color-coded.

Phylogeny:

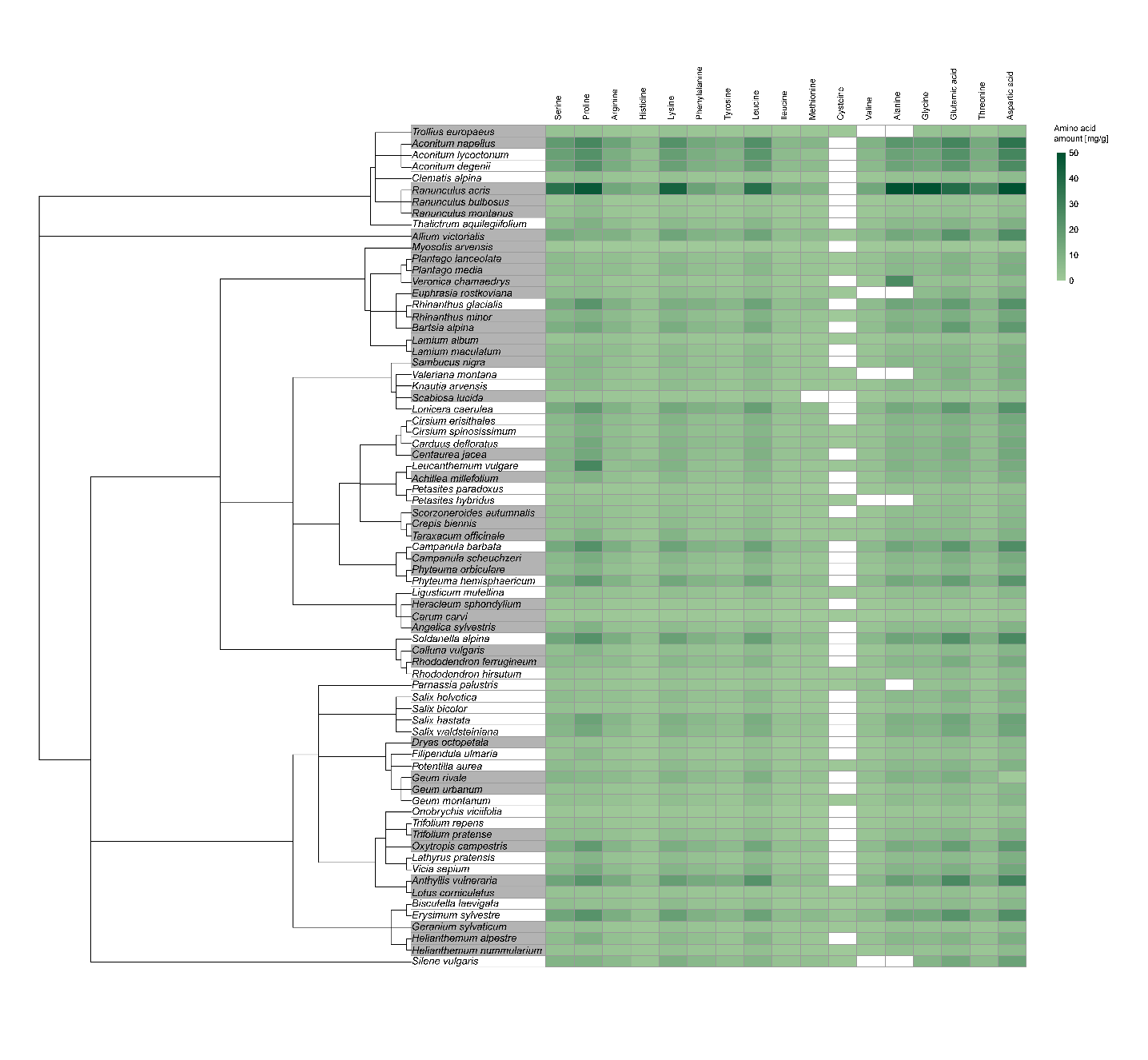

Figure S4: Pollen AA profiles of plant species. AAs are shown at the top with different concentrations depicted by different shades of green. Phylogenetic relationships are given on the left. Plant species with a grey background were collected by pollinators, plant species with a white background were not collected.

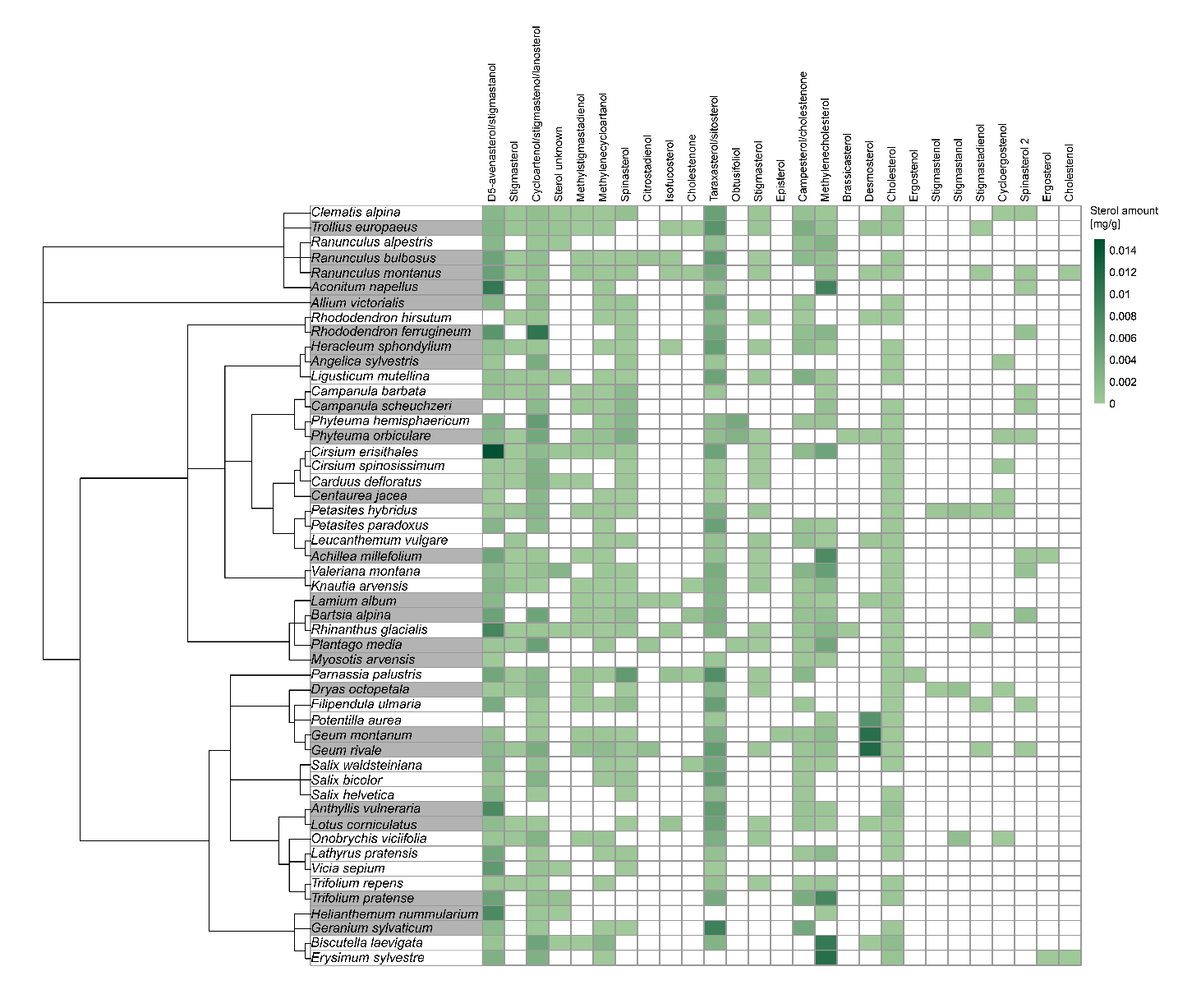

Figure S5: Pollen sterol profiles of plant species. Sterols are shown at the top, with different concentrations depicted by shades of green. Phylogenetic relationships are given on the left. Plant species with a grey background were collected by pollinators, plant species with a white background were not collected.
